# Generalized prefrontal task-progress sequences efficiently encode task structure

**DOI:** 10.64898/2026.10.20.760165

**Authors:** Ousama Tarahi, Kevin Koshy, Jonas-Frederic Sauer

**Affiliations:** Centre for Integrative Physiology and Molecular Medicine, Dept. Systemic Neurophysiology, Medical Faculty, Saarland University, Homburg, Germany

## Abstract

The medial prefrontal cortex (mPFC) occupies a central role in executive function including decision-making, rule learning and cognitive flexibility. Here, we investigated how mPFC populations represent the animal’s progression along sequential task stages. Freely moving mice performed an odour-guided two-choice task requiring sensory sampling, action selection, navigation, and reward collection. Population activity in the mPFC formed sequences that evolved through cue sampling, movement towards the reward sites, and reward-related epochs. These sequences exhibited a shared temporal organization across trial types, pointing to a generalized encoding of task progress by prefrontal neurons. Artificial neural networks trained on the same task developed task-progress sequences, and experimentally disrupting task-progress tuning in these networks impaired learning efficiency. Our results indicate that mPFC activity organizes behavioural episodes along correlated trajectories and suggest that task-progress tuning offers an efficient solution for the encoding of task space during multi-stage behaviour.

## Introduction

Goal-directed behaviour requires neural circuits to integrate multiple information streams, including sensory cues, the agent’s position within the task environment, and the location of potential targets. In many cases, such behaviours do not reflect instantaneous stimulus– response mappings but instead evolve in a sequential manner. For instance, in cued decision making tasks, the agent needs to sample an informative cue, select the appropriate reward target from memory, and plan and execute a correct trajectory to the current target. The multi-stage nature of such tasks raises the question how brain circuits integrate these diverse aspects. The medial prefrontal cortex (mPFC) is a crucial candidate brain region to implement this computation, given its central role in spatial working memory, rule learning and navigation^1–3^. Neurons in the mPFC have been shown to encode learned stimuli^4–6^, choices^7,8^, and the space along which the animal navigates^9^, often in a mixed-selective manner^7,10^. However, it is not fully understood how mPFC populations encode the seamless transition through the various stages of such sequentially structured tasks.

A prominent hypothesis is that the mPFC constructs cognitive maps of abstract task structure. Prefrontal and orbitofrontal regions have indeed been proposed to represent latent task states that must be inferred from the structure of the task^11–13^. Moreover, recent work in an ABCD task^14^, in which mice learned to cover abstract spatial sequences to obtain a reward, demonstrated that neural activity can reflect an animal’s position within a structured sequence of states rather than in physical space. These findings suggest that prefrontal neurons reflect task-progress tuning, meaning that individual neurons show activity peaks relative to the agent’s progression towards a target^14,15^. In the spatial domain, this conclusion is supported by similar tuning of mPFC cells in a maze in which paths of varying length can be chosen by the animal^16^. However, it remains to be tested whether task-progress representations are the preferred mode of operation of prefrontal circuits, in particular when the animals must bridge sensory sampling, action selection, and outcome processing within a single trial.

Here, we address this question in an odour-guided two-choice task performed by freely moving mice^17^. The task naturally comprises distinct behavioural stages within each trial, including odour sampling, movement away from the odour port, and arrival at the reward site, making it well suited for examining whether mPFC activity is organized around a latent variable that spans the full trial. Rather than focusing on whether a single epoch contains task information, we asked whether population activity follows a structure that tracks progression through the individual behavioural epochs within a trial. To examine whether such a representation could arise from task demands alone, we further trained recurrent neural networks on the same task and asked whether they develop internal dynamics that mirror the structure observed in mPFC.

We found that mPFC activity is organized into task-progress sequences that extend from odour presentation through movement and into reward-related epochs. Task-progress sequences display a shared structure between trial types while maintaining enough differences to allow predicting the current trial type from sequential information alone. In other words, mPFC activity along separate trajectories towards distinct targets remains correlated but discriminable. Similar task-progress-like dynamics emerged in recurrent networks trained on the task, and disrupting task-progress organization in these artificial networks impaired learning efficiency. Together, these findings suggest that task-progress tuning may provide an efficient way to organize prefrontal activity across an unfolding behavioural episode.

## Results

### Prefrontal task progress *sequences* during olfaction-guided *decision* making

To probe sequential neuronal activity in the mPFC, we analysed previously published 1-photon calcium imaging data from mice that executed an olfaction-guided decision task, in which one (of two possible) odour cues indicated whether to approach the left or right arm of an M-maze^17^. In this task, the animals start a trial by sampling the odour, move subsequently through the outward arm, and enter the reward site to receive a reward. Thus, each trial is composed of a series of intermingled non-locomoting (sampling, reward consumption) and movement steps (outward motion, **Fig. 1A**). Previous analyses of this^17,18^ and related tasks^19–21^ revealed spatial tuning of mPFC pyramidal cells as the animals moved towards the target. In such analyses, periods of movements are extracted to then assess the spatial tuning functions of mPFC neurons. Here, we took the inverse approach and searched for repeating sequential activity patterns without pre-selecting specific task epochs. First, we applied RasterMap^22^, an algorithm for pattern detection. RasterMap identified repeating sequential activity patterns starting during the sampling period and extending through the reward epoch, thus covering both periods of movement and immobility (**Supplementary Fig. 1**). To formalize this finding, we developed a sequence detection method in which the calcium activity of prefrontal pyramidal cells during a single trial (starting from entry into the odour sampling zone) served as a template and subsequent trials were scanned for matching sequential activity (**Fig. 1A**). Importantly, the detection method was applied entirely in the time domain, thus ignoring whether or not the animal actively moved through space at any given time point. Significant matches were identified for time points at which the deviation in the time-to-peak distribution of the participating neurons remained below the 0.001th percentile of that computed from randomly permuted distributions (Methods). As a control, we performed sequence detection on surrogate data created by randomly time-shifting the calcium activity of each neuron before sequence detection, thereby removing any correlated temporal structure. Focussing on the odour presentation of a given trial as the starting point for the template, we detected matching sequences throughout the session (**Fig. 1B**). Both the proportion of trials with significant matches (reactivation proportion) and the trial-wise template-match similarity (Pearson’s r of time-to-peak distributions) indicated a significant effect over control data (**Fig. 1C**). Importantly, although our algorithm did not constrain when during a trial potential matches could be found, they were significantly enriched around trial starts, suggesting that sequences reoccur tied with odour sampling (**Fig. 1D**). Running the detection analysis with randomly chosen template start times instead, we observed a significant drop in reactivation proportion and template-match similarity as the starting point moved away from the sampling period (**Supplementary Fig. 1**). The detected sequences spanned from the sampling epoch through the outgoing trajectory and beyond arrival at the reward site. While more neurons contributed to sequences during early sampling, likely reflecting cue-evoked responses^5^, we found no difference in the number of neurons participating in sequence activity during mobile and immobile task phases, suggesting that sequential activity seamlessly evolves through non-locomoting and movement epochs (**Supplementary Fig. 2**).

**Figure 1:**
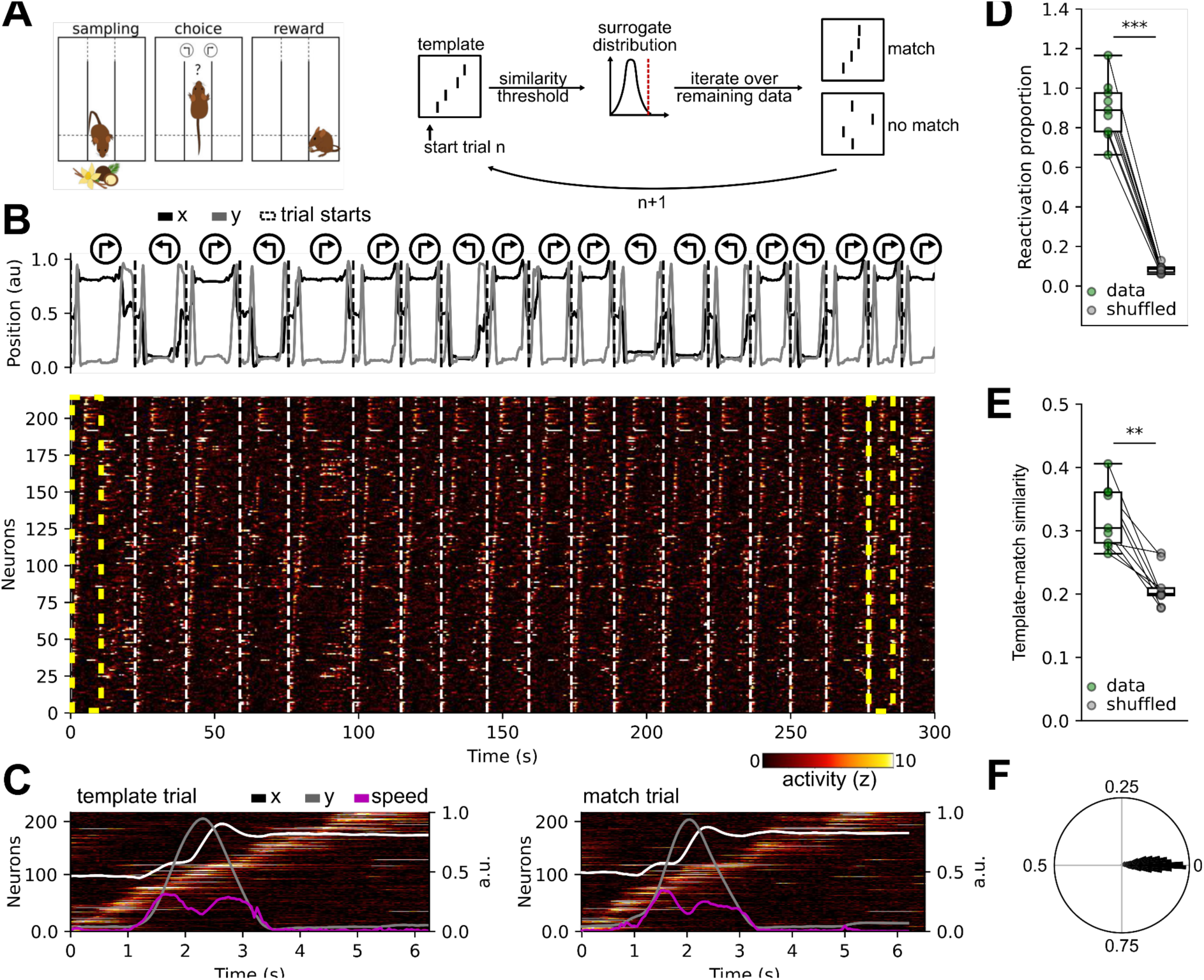
Prefrontal task-progress tuning in a decision making task. **A** Left: Schematic of the task (adopted from Muysers et al^17^ under CC BY 4.0, https://creativecommons.org/licenses/by/4.0/). Mice are presented in each trial with one of two odours (vanilla/coconut), each indicative of a reward site in one of the side arms. Right: Schematic of the procedure to identify neuronal sequences from 1-photon calcium imaging data. The activity of prefrontal neurons (symbolized by vertical ticks, the actual analysis was performed on normalized calcium traces) is sorted by the time of peak relative to entry into the sampling zone (trial start). Iterating over the remaining data of the session, the algorithm finds recurrences of the same sequence (matches) identified by a similarity criterion obtained against shuffled data (see Methods for details). Note that the procedure iterates over template trials such that each trial serves as a template once. **B** Top: example behavioural data (x- and y-coordinates of the mouse) of one session. Symbols on top indicate whether each trial is left- or rightward going. Dashed lines indicate trial starts (i.e., entry into the odour sampling zone). Bottom: Neuronal activity during the same session, sorted by time-of-peak of the first trial, reveals recurring sequential activity throughout the session. Boxed areas (left: current template, right: example match) are shown at larger magnification in C. **C** Left: Neuronal activity sorted by peak activity with overlaid x- and y-coordinates as well as the speed profile of the template trial (given in arbitrary units). Right: An example match trial corresponding to the right boxed area in B, in the same neural sorting as the template trial. Identified sequences progress through periods of both movement and immobility. **D** Quantification of the proportion of trials with significant reactivation of the template sequence. p=8.5e-8, t=18.2, paired t-test. **E** Quantification of the template-match similarity (Pearson’s r of time-to-peak distributions). p=0.004, W=0, Wilcoxon signed rank test. **F** Number of detected matches for all animals (n=7773) plotted as a function of relative task time ranging from trial start (0) to the return of the animal to the sampling site (1). Detected sequences are enriched around trial start. All data based on n=9 mice. Circles in D, E show mouse averages, boxes show median and first and third quartiles.

To corroborate the observation that neuronal sequences consistently evolved across immobile task epochs, we trained decoding models to predict the elapsed time from the beginning of the sampling and reward period, respectively (i.e., covering the initial phase of both non-moving trial epochs). Decoders trained on the data significantly outperformed those trained on shuffled data (**Supplementary Fig. 3**). Extending upon previous accounts of ordered activity of prefrontal neurons during active movement, these data jointly suggest that task-progress sequences extend from odour sampling through target-directed motion until after arrival at the reward site.

### Generalized encoding of task progress

Movement-related prefrontal sequences display a generalized representation of task space, meaning that spatially distinct trajectories are reflected in similar (albeit not identical) sequences^15,18^. To test whether such generalized representations hold true for task-progress sequences identified here, we separately analysed for any given template the matches for the same (i.e., template and match are both either left or right trials) or opposite trial type (i.e., template: left, match: right, or vice versa). Same sequences showed significantly larger template-match similarity than opposite sequences. However, in quantitative terms this difference was small (0.34 ± 0.02 vs. 0.31 ± 0.02 (mean ± sem)), suggesting a similar sequence structure for both trial types (**Fig. 2A**). To further corroborate that finding, we expressed the activity of the neurons as a function of task phase ranging from the of entry into the odour sampling zone (time point zero) to 500 ms after arrival at the reward site (time point one; **Fig. 2B**). This time span was chosen to match sequence duration and to cover both immobile and movement periods. We computed average phase tuning functions separately for left- and right-going trials, respectively. Correlating average phase tuning functions between both trial types (termed phase similarity) averaged at 0.45 ± 0.03, suggesting substantial similarity in phase tuning. As a comparison, we calculated phase tuning functions for the first and second half of the session, separately for left- and right-going trials (phase consistency). Phase consistency averaged at 0.61 ± 0.03, therefore close to (albeit significantly larger than) the phase similarity reported above (**Fig. 2C**). Similarly, phase similarity computed for the odour sampling phase only (i.e., for the task epoch from trial start to the onset of the outward trajectory) was in a similar range as the phase consistency of the same task epoch (**Fig. 2D**). These results jointly suggest that neuronal sequences during both trial types are correlated in structure.

**Figure 2:**
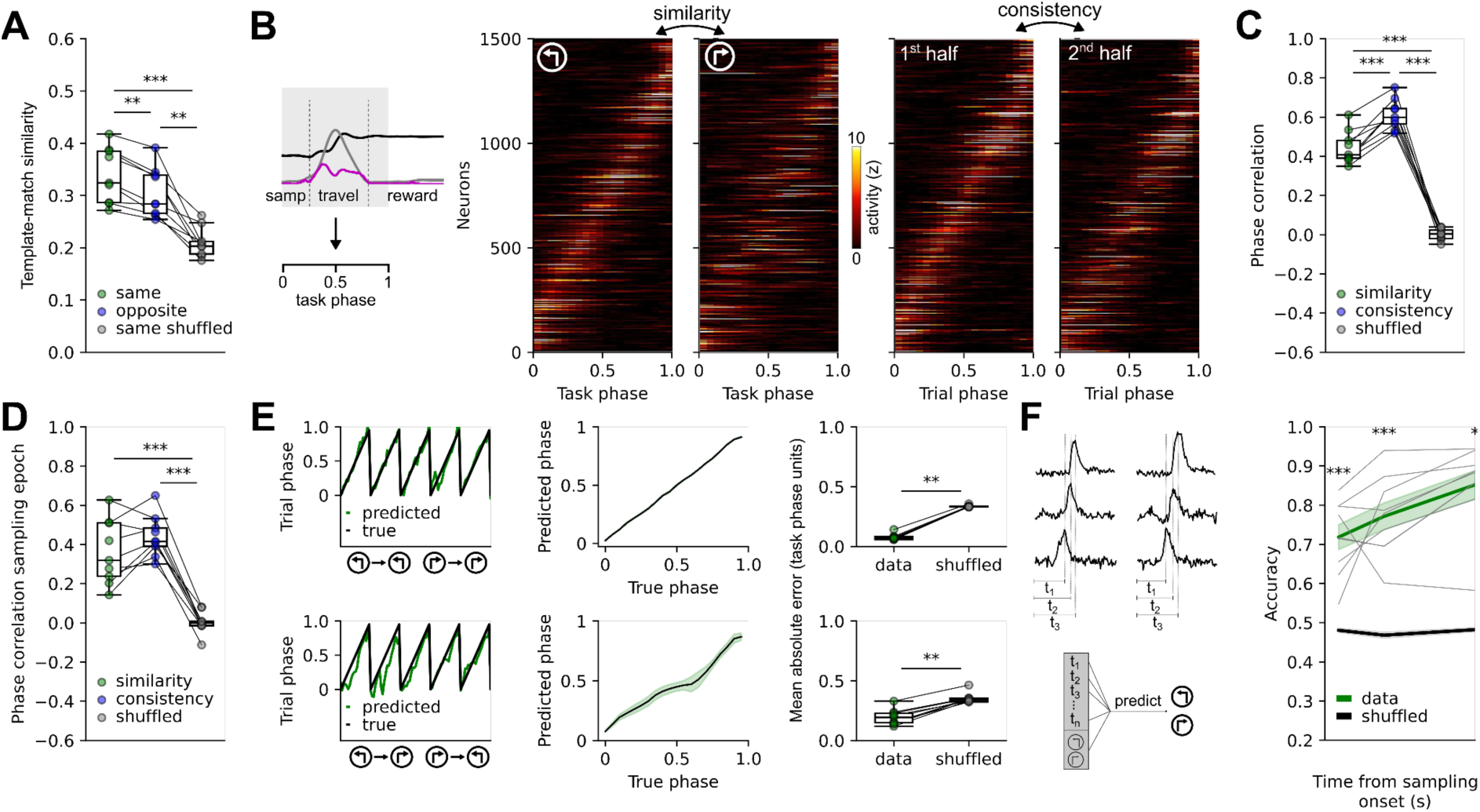
Task-progress sequences show generalized properties. **A** Average template-match similarity for template-match pairs of the same trial type (both left or both right, ‘same’) exceeds that of templates and matches coming from different trial types (‘opposite’, p=0.001, t=5.87) and of shuffled same data (p=0.0005, t=6.68; opposite vs. shuffled: p=0.001, t=5.89). One-way repeated measures ANOVA followed by paired t-tests with Šidák correction. **B** Task-phase analysis. Left: The time from the start of odour sampling to 500 ms after arrival at the reward site was converted to a task phase ranging from 0 to 1. Grey area illustrates the converted period of the example trial. Middle: Average activity of all neurons as a function of task phase sorted by peak phase for left-going trials. The activity is shown in the same sorting for right-going trials. Similarity between both populations of tuning functions was measured with Pearson’s r. Right: Same as middle but for the first half vs. the second half of the session to measure within-session consistency (only left-going trials are shown). **C** Average similarity is lower than consistency (p=0.0002, t=-7.3) but significantly different from chance estimated from consistency values for shuffled data (p=2e-6, t=13.9; consistency vs. shuffled: p=e-7, t=19.5). One-way repeated measures ANOVAs followed by paired t-tests with Šidák correction. **D** Same as C but with only the task phase during the sampling phase considered (0: entry into sampling area, 1: exit from sampling area). There is no difference between similarity and consistency (p=0.059, t=-2.2) with both being significantly above chance level (similarity: p=0.0002, t=6.5, consistency: p=3e-7, t=15.1). One-way repeated measures ANOVAs followed by paired t-tests with Šidák correction. **E** Decoding task phase from calcium activity. Top row: same trial decoding (training & testing on either left- or right-going trials). From left to right: example true (black) and predicted task phase (green) of a linear regression model, summed predicted phases over all mice, decoding error for data vs. phase-shuffled data (p=0.004, W=0). Bottom row: same as top row but with only the task phase during the sampling phase considered. p=0.004, W=0. Wilcoxon signed-rank tests. **F** Decoding trial identity (i.e., left- vs. right-going) based on temporal information. A logistic regression model was trained on time-to-peak signals of mPFC neurons to predict trial identity. Right: Decoder accuracy exceeds random controls obtained from shuffled trial labels (grey) for different time windows considered for decoding (data vs. shuffled: p=1.7e-6, F=152, effect of time window: p=0.044, F= 5.0, interaction effect: p=0.03, F=5.7, two-way repeated measures ANOVA; post hoc comparisons for 1, 2 and 4 s time window: p=0.0002, 6e-5, 0.012, t=7.4, 8.9, W=0, paired t- and Wilcoxon signed rank tests with Šidák correction. All data based on n=9 mice. Circles in A, C, D, E show mouse averages, boxes show median and first and third quartiles. Thick lines in E, F show mean, grey lines individual mice, shaded areas sem.

To further investigate the correlated sequence structure, we trained decoding models to predict the current task phase from calcium activity. Within each trial type (i.e., training and testing on either left- or right-going trials), the model significantly predicted the task phase (**Fig. 2E**). When we performed cross-trial decoding (training on left-going trials, testing on right-going trials or vice versa), we moreover observed significant task phase decoding, reinforcing our above conclusion about the generalized nature of task-progress sequences (**Fig. 2E**). Finally, given the correlated sequence structure for both trial types, we tested whether the information contained in the sequential activation of neurons is sufficient to predict the trial outcome from the order of activity alone. To test this, we trained a decoder with the time of peak activation of each neuron and trial labels. For a range of times from trial onset used for decoding, the decoder significantly predicted trial outcome versus shuffled controls (**Fig. 2F**). Taken together, these data suggest that prefrontal activity contains generalized information about task progress while maintaining trial type decodability.

### Generalized task progress sequences emerge in an artificial network

Generalized task-progress sequences might reflect an efficient mechanism to simultaneously encode both task progress and current trial type. We therefore hypothesized that this property would be ideally suited for task learning. To test this proposal, we investigated task-progress tuning in artificial neuronal networks, which enables us to experimentally alter the network’s capacity to form task-progress tuning. To model the behavioural task, we trained a closed-loop recurrent rate network (N = 200, connection probability p = 0.4) to predict *K* fixed two-dimensional trajectories *o_k_(t)* composed of x- and y-coordinates from a single graded cue that represents the odour identity (**Fig. 3A**):

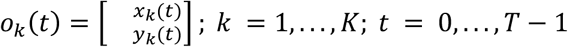

**Figure 3.**
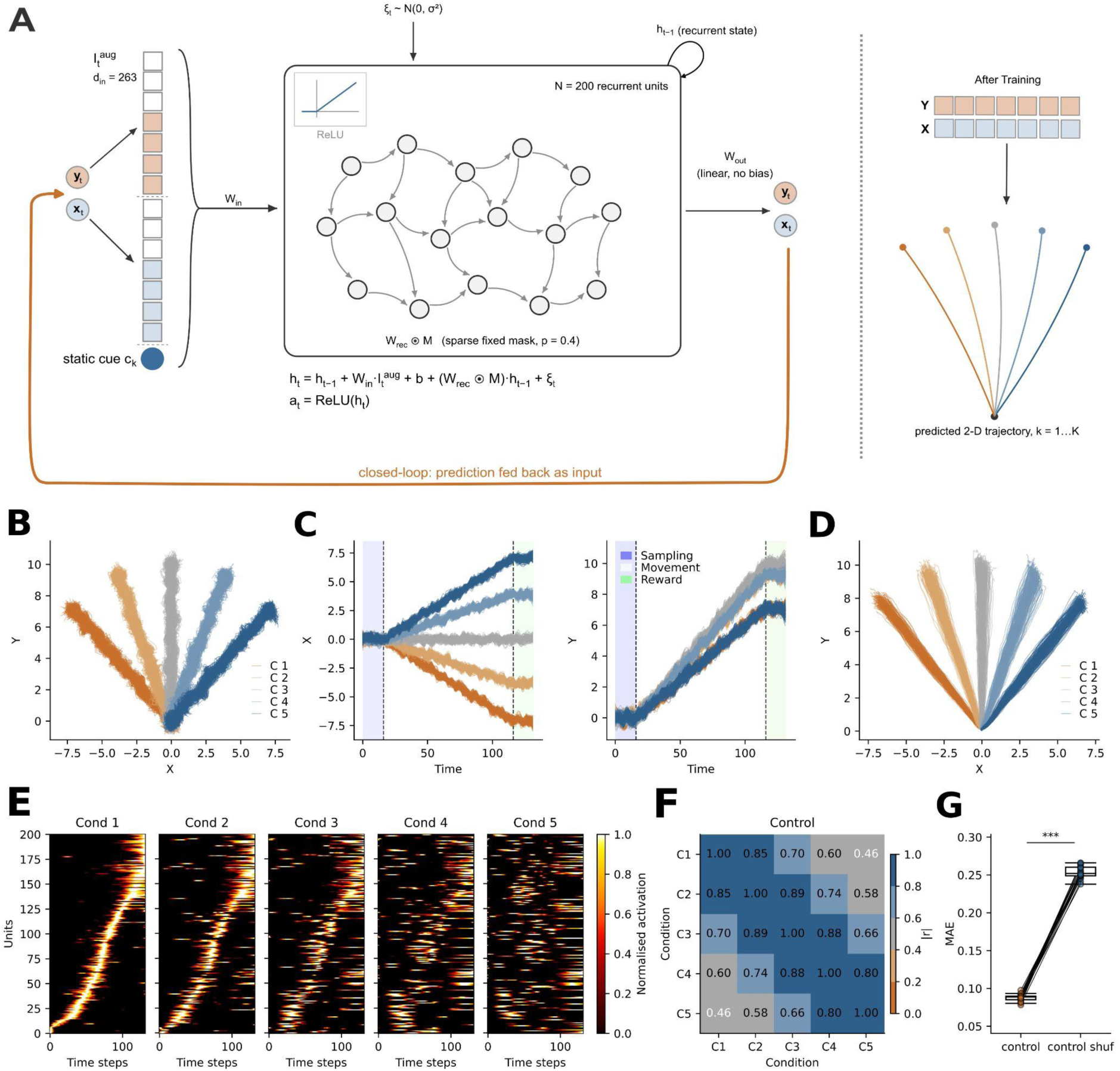
A closed-loop recurrent network spontaneously develops generalized task-progress sequences. **A** Schematic of the closed-loop recurrent rate network. A static scalar cue c encodes trajectory identity and is concatenated at every time step with the network’s own previous 2-D output; the recurrent layer (N = 200 units, connection probability p = 0.4, fixed sparse mask) drives a bias-free linear readout of the behavioural coordinates. No teacher forcing is used at any point — the rollout is free-running from the first to the last step. **B** Target output trajectories for the K = 5 trained conditions (C1-5) in the behavioral (X–Y) plane, colour-coded by condition. Each trajectory comprises a stationary sampling phase, a condition-specific curved movement phase, and a stationary reward phase. **C** Temporal evolution of the X (top) and Y (bottom) coordinates of the same target trajectories. Background shading denotes the sampling (blue), movement (white), and reward (green) epochs. **D** Network-generated (predicted) trajectories after training, shown in the same 2D behavioral (X–Y) plane. **E** Trial-averaged hidden-unit activation heatmaps for each condition for the control network. Units are sorted by peak-activation latency, with the ordering fixed from Condition 1. A consistent diagonal band is visible across all five conditions, indicating a shared, sequential temporal scaffold reused across conditions. **F** Representational similarity matrix, from a representative control network, showing the mean pairwise absolute Pearson correlation between condition-averaged hidden-unit tuning curves. Values indicate the magnitude of representational similarity, irrespective of the sign of the correlation. mean cross-condition (|r|) = 0.72 ± 0.01 (mean ± SD). **G** Cross-condition decoding of task phase: a linear regression model trained on condition 1 and tested on the remaining conditions achieves substantially lower mean absolute error than a label-shuffled control (t = −64.89, p = 2.47e-13, paired t-test), in which phase labels are randomly permuted across training samples. This demonstrates that the temporal scaffold generalizes across conditions.

Since our model is not constrained by physical task space, we chose *K=5* trajectories (or conditions) for the main analyses (**Fig. 3B**). The target trajectories consisted of three behavioural phases: a stationary cue sampling phase, a condition-specific movement phase, and a stationary reward phase (**Fig. 3C**), thus closely mimicking the overall structure of the task as performed by the animals. The network was solely optimized to reproduce the behavioural output *o_k_* (**Fig. 3D**), without constraints. Upon training, the hidden units became active in a highly organized, cue-locked temporal sequence throughout the trial. Sorting units by their peak activation latency revealed a diagonal-band structure, similar to the one observed in prefrontal recordings (**Fig 3E**). Furthermore, condition-specific tuning curves showed high representational similarity across behavioural conditions (mean pairwise |r| = 0.72 ± 0.01, mean ± SD; **Fig. 3F**). Finally, training a decoding model on one condition allowed significant prediction of task time in other conditions (**Fig. 3G**). These results indicate that the network spontaneously developed a shared temporal scaffold that is reused across multiple trajectories, resembling generalized task-progress tuning as observed for mPFC pyramidal neurons.

### Generalized task progress sequences enable efficient learning in artificial networks

To establish conditions under which the artificial network is forced to solve the task without resorting to generalized task-progress tuning, we added a differentiable penalty on the absolute cross-condition correlation of per-unit tuning curves, weighted by α>0. This additional loss term constrains hidden units when they exhibit similar activity patterns across different behavioural conditions *K*, hence pushing each condition to develop more distinct neural representations and penalising the emergence of generalized task-progress tuning (**Fig. 4A**). Importantly, the penalty selectively modified the organization of network dynamics without changing the underlying architecture or connectivity statistics: the distribution of active recurrent weights in constrained networks was statistically equivalent to that of seed-matched controls within a pre-specified margin of ±0.00063 (TOST, p = 2.02e-20), and individual connection weights remained strongly correlated between the two (Pearson r = 0.799; **Supplementary Fig. 4A**). Therefore, differences in learning dynamics and representational structure between the two networks are unlikely to be explained by changes in network capacity or connectivity, and instead point to the loss of shared coding.

**Figure 4.**
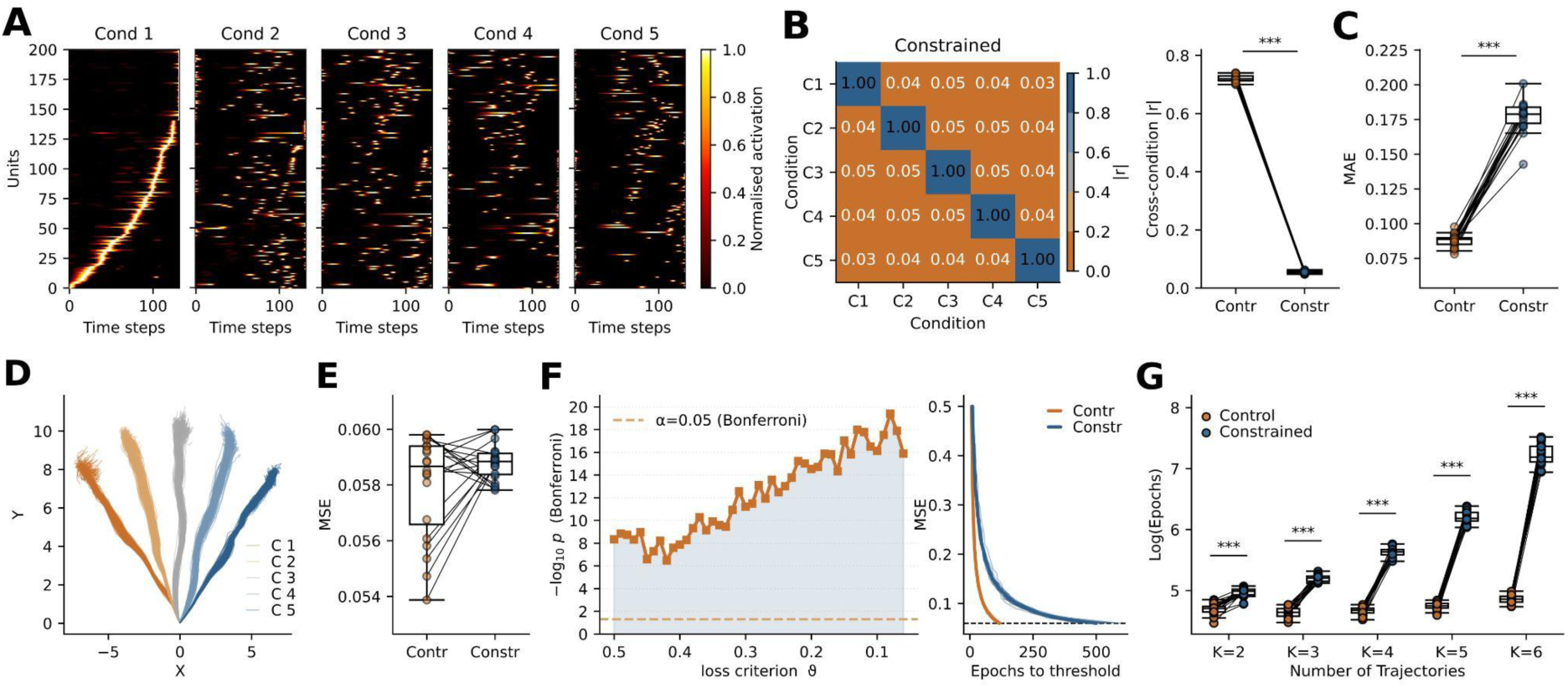
Suppressing generalized task-progress coding preserves asymptotic performance but impairs learning efficiency, in proportion to task complexity. **A** Trial-averaged hidden-unit activation heatmaps for each condition for the constrained network (K = 5, α = 1), same sorting convention as Fig. 3E. Sequential structure is preserved only for the condition used to define the sort order (condition 1), while absent for the remaining conditions, reflecting condition-specific rather than shared coding. **B** Loss of cross-condition representational similarity in the constrained network. Left, representational similarity matrix for constrained network, showing reduced similarity across conditions (cf. Fig. 3F for control). Right, mean absolute cross-condition correlation of hidden-unit tuning curves, decreasing from 0.72 ± 0.01 in the control network to 0.05 ± 0.01 in the constrained network (t=207.7, p= 2.36e-33, paired t-test, n=20). **C** Comparison of cross-condition decoding of task phase. A linear regression model trained on Condition 1 and tested on the remaining conditions achieved substantially lower mean absolute error when applied to data generated by the control network than when applied to data generated by the constrained network (t= −32.58, p= 3.89e-18, paired t-test, n=20). **D** Generated output trajectories of the constrained network for all five conditions, shown in the same 2D behavioral (X–Y) plane (cf. Fig. 3D for control). **E** Final task performance of control and constrained networks. Final post-convergence task MSE for control and constrained networks, showing statistically indistinguishable asymptotic behavioral performance (TOST equivalence test, paired, bound = ±0.006 [10% of θ*], dof = 19, p =4.2e-10). **F Left**, Bonferroni-corrected significance (–log₁₀ p) of the control-vs-constrained difference in epochs- to-criterion, computed across 45 loss criteria spanning θ (paired t-tests, Bonferroni-corrected, dof = 19). The difference was significant at every criterion tested (Bonferroni-corrected p range: [4e-20, 3.33e-7]; |t| range: [9.79, 50.9]), with larger separations at stricter thresholds. **Right,** task MSE as a function of training epoch for control and constrained networks; dashed line indicates the convergence threshold (θ* = 0.06). **G** Convergence speed across number of trajectories K for both networks. Number of training epochs required for control and constrained networks to reach the predefined behavioral loss criterion (θ*=0.06) as a function of K. Constrained networks required significantly more epochs than their seed-matched controls at every K (paired t-tests, dof = 19, n = 20 seeds: K = 2, t = −11.59, p = 4.6e-10; K = 3, t = −24.79, p = 6.2e-16; K = 4, t = −36.79, p = 4.0e-19; K = 5, t = −33.12, p = 2.9e-18; K = 6, t = −22.68, p = 3.2e-15). The training gap widens systematically with task complexity.

Constrained networks exhibited a marked reduction in cross-condition similarity of hidden-unit tuning curves, with mean absolute correlation decreasing from 0.72 ± 0.01 in control networks to 0.05 ± 0.01 in constrained networks (mean ± SD; *t* = 207.7, *p* = 2.36e-33, paired *t*-test; **Fig. 4B**). This loss of cross-condition similarity indicates that the constrained network developed more distinct condition-specific representations, in contrast to the shared representational structure observed in control networks (**Fig. 4B**; cf. **Fig. 3F**). In line with the reduced correlation, decoding analysis applied to constrained networks revealed substantially reduced across-condition task-progress decodability (**Fig. 4C**). The control networks concentrated their activity within a compact low-dimensional subspace, while constrained networks distributed their activity across a larger number of dimensions, offering more condition-specific neural codes (**Supplementary Fig. 4B**). Expectedly, suppressing this shared structure was accompanied by an increase in dimensionality. Both the participation ratio and the dimensionality required to explain 90% of the variance increased significantly with the introduction of penalty (**Supplementary Fig. 4C**). As a result of the larger dimensionality, constrained networks operated at significantly lower mean hidden-unit activity than their seed-matched controls (**Supplementary Fig. 4D**).

Both control and constrained networks solved the task, converging to the same asymptotic behavioural loss (**Fig 4D,E**). However, the constrained network required significantly more training epochs before reaching a behavioural criterion *E\** (epochs-to- criterion, 115.5 ± 7.2 vs 500.1 ± 53.2 epochs, with a mean difference of 384.6 epochs and a percentage increase of 333.1%; paired t-test: |t|=33.12, p=2.9e-18, dof=19; **Fig. 4F**). This effect was robust across different loss criteria, with the difference in epochs-to-criterion remaining significant at every threshold tested and becoming larger at stricter criteria (**Fig. 4F**). We found control networks to robustly outperform constrained models in terms of learning efficiency for varying data sizes and trajectory geometries (**Supplementary Fig. 5A-B**). Moreover, modulating the penalty strength term α identified a dose-dependent effect on learning efficiency (**Supplementary Fig. 5C**). Finally, control simulation with activity-penalized networks indicated that the reduced mean activity observed in constrained networks could not explain their lower learning efficiency (**Supplementary Fig. 6**). Shared representations thus improve optimization efficiency while preserving final performance.

To exclude the possibility that these results are contingent on the specific architecture of our model, we additionally trained classical recurrent neural networks on the task. With this approach, we again observed generalized task-progress sequences in the control but not the penalized model. Similar to our closed-loop network approach, we found that penalized models required significantly more training epochs to learn the task (**Supplementary Fig. 7**). Taken together, these results suggest that the occurrence of task-progress sequences enables the network to learn the task efficiently.

### The learning advantage of generalized task-progress encoding scales with trajectory number

If the benefit in learning stems from reusing a common temporal scaffold across conditions, its magnitude should scale with the number of conditions. To explicitly test this prediction, we ran independent simulations with an increasing number of conditions *K.* In the simplest version of the task with two conditions, the constrained network converged only slightly slower than the control network, suggesting that the constrained network could still learn distinct representations for each condition with little additional optimization cost (**Fig. 4G**). However, as task complexity increased with the number of trajectories *K*, the constrained network required substantially more training epochs to converge than the control network, resulting in an increasing training gap (**Fig. 4G**). This learning disadvantage was robust, as a two-way repeated-measures ANOVA revealed a strong main effect of network type (control vs. constrained; F = 2677.1, p = 6.52e-22). Furthermore, a significant network × criterion interaction (F = 1069.01, p = 1.48e-36) indicated that the effect of network type varied as a function of performance criterion, increasing at more stringent thresholds. Together, these results demonstrate that the penalty selectively affected learning dynamics, producing distinct performance trajectories between control and constrained networks rather than simply causing a uniform shift in performance.

## Discussion

We identified task-progress sequences in a 2-choice olfaction-guided decision making task. These sequences span across behavioural epochs of immobility and movement. They display generalized features in the form of correlated sequence structure across trajectory types while maintaining sufficient distinctiveness to allow trial type decodability.

Since prefrontal task-progress sequences are not explicitly linked with motion, they extend beyond a purely spatial^1,19–21,23,24^ or purely temporal representation^25–27^. Rather, they are in line with the more general notion of sequential activity motifs as a central coding element to reflect external structure. For instance, during visuo-spatial working memory in monkeys, prefrontal sequences span from cue presentation through working memory delay to virtual navigation^28^. Moreover, task-progress tuning in the *ABCD* task^14^, in which progression towards the current goal is more prominently reflected in prefrontal activity than current location, are in line with the task-progress sequences reported here. Together with recent observations of task-progress tuning across spatially distinct paths^16^, these findings suggest that generalized task-progress tuning is a widely used encoding paradigm of prefrontal circuits.

Task-progress sequences are likely formed during task learning via the strengthening of incoming and recurrent synaptic connections. This might create preferred and consistent activation paths through the network, in which activity propagates from cue-selective starter neurons that are initially activated by an incoming input towards subsequent neurons. Such ordered activity is a recurring motif in neuronal networks. For example, driving activity in a single pyramidal cell induces a sequence of activity in the turtle cortical network^29^. Moreover, neocortical neurons in the mouse follow a stereotype activation sequence during UP-state onset^30^. Finally, during time interval estimation, prefrontal time cells activate sequentially^31^. A central open question is how the propagation of activity is precisely controlled in the network: During immobility, arriving sensory cues are expected to change less over time compared to when the animal is actively traversing the arena. This is because movement is accompanied by constantly changing visual and tactile input. As a result, the propagation of sequential activity might be driven more by internal circuit dynamics during immobility and less by external input, with the opposite ratio during movement. Neuromodulatory state, for instance an increase in acetylcholine release during movement compared to rest^32–34^, might be a candidate mechanism supporting such a network transition.

In our artificial networks, task-progress sequences naturally emerge, suggesting that this form of representation is not incidental but arises from task demands. The artificial network exhibits an activity organization similar to that observed in the biological population, in which a substantial component of the response is condition-invariant. Consequently, population dimensionality is low because a common sequential representation is shared across conditions and captures most of the variance. In contrast, constrained networks that develop condition-specific representations must recruit additional dimensions, producing a higher-dimensional population structure (**Supplementary Fig. 4C**). This came at the cost of lower overall activity levels, consistent with control networks concentrating a large, shared signal in a compact low-dimensional code while constrained networks spread comparatively weaker activity across a larger number of dimensions. Importantly, reduced neural activity alone did not explain why constrained networks learned more slowly. Even when their activity was restricted to half that of the control networks, they still maintained the shared task-progress representation and learned at least as quickly as the controls (**Supplementary Fig. 6**).

The shared task-progress code in our models is not necessary for correct behaviour since both control and constrained models show indistinguishable asymptotic performance. This is because the recurrent architecture is likely expressive enough to reach the same performance with condition-specific representations. Yet, the control network converges rapidly, while the constrained network requires a longer optimization trajectory to arrive at the same solution. The control network likely accelerates learning by reusing a temporal scaffold across conditions, whereas the constrained network must learn more independent solutions due to reduced representational sharing. Consequently, shared representations improve optimization efficiency while preserving final performance.

Our results have a well-established counterpart in machine learning, where learning one task helps the model learn another related task by leveraging shared useful information^35,36^. Additionally, parameter sharing is essential for efficient learning by enabling related tasks to reuse common internal representations rather than learning independent solutions^37^. In our recurrent network, this principle is reflected not only in shared parameters but also in the emergence of common sequential activity patterns across conditions, suggesting that related tasks can be solved by reusing a shared dynamical structure. Learning related tasks leads to the emergence of a low-dimensional neural schema that facilitates subsequent learning by limiting the extent of the required weight changes^38^. Our contribution examines the sharing of a dynamical trajectory across conditions, and directly alters that sharing while architecture, connectivity, targets and behavioral loss are held fixed. Consistent with the mentioned studies, our results suggest that shared recurrent dynamics provide a similar computational advantage, enabling more efficient optimization without altering the final achievable performance.

While any task involving transitions through a sequence of states is likely to invoke sequential neural representations, we hypothesize that the emergence of generalized task-progress tuning has been evolutionarily favoured due to the computational efficiency of this solution. This feature of prefrontal circuits is moreover well aligned with the circuit’s role in generalizing learned experiences to novel environments^39^.

### Limitations of our study

Our findings should be interpreted in light of several limitations. First, the network model is optimized using gradient descent with supervised trajectory matching that is not intended to model the biological learning mechanisms of the brain. Accordingly, our conclusions concern the computational consequences of shared versus separated neural representations and the resulting optimization landscape, rather than the synaptic plasticity rules by which animals acquire these representations. Second, the correlation penalty serves as an experimental intervention rather than a biologically grounded mechanism. While it enables us to test the computational role of generalized tuning by selectively restricting representational sharing, it does not identify the circuit or plasticity mechanisms that would produce such constraints in vivo. Moreover, because the penalty is applied to the absolute correlation between condition-averaged tuning curves, it imposes a stricter constraint than the signed similarity measured experimentally, excluding both positively and negatively correlated forms of representational reuse. Finally, control networks achieved a cross-condition correlation of 0.72 ± 0.01, whereas mPFC phase tuning exhibited a correlation of 0.45 ± 0.03 across trial types. Thus, the shared structure observed in cortex is weaker than that emerging in networks optimized solely for behavioral accuracy (but see our analysis of cross-correlation penalties of different strength in **Supplementary Fig. 5**), suggesting that the artificial network finds a solution that is more generalized than the one implemented in the mPFC.

## Acknowledgements

We thank Hanna Muysers for the previous acquisition of the dataset used in this study, Daniel Rein for technical assistance, and Davide Gobbo for input on this work. This work was supported by Deutsche Forschungsgemeinschaft (grants JSA 3609/2-1, JSA 3609/2-2 (FOR5159) and JSA 3609/3-1 (CRC/TRR384 IN-CODE) to J.F.S.. This work is supported by ERC grant DRIFT-rules (DLV-101229453). Funded by the European Union. Views and opinions expressed are however those of the author(s) only and do not necessarily reflect those of the European Union or the European Research Council Executive Agency. Neither the European Union nor the granting authority can be held responsible for them.

## Author contributions

OT designed, implemented and analyzed closed-loop recurrent rate networks. KK and OT designed, implemented and analysed recurrent networks. JFS analysed neural data. OT and JFS wrote the original manuscript. All authors revised and edited the manuscript.

## Data and code accessibility

The experimental data used in this study was previously published^17^ and can be found at https://doi.org/10.5281/zenodo.10528244. Code for the analysis of experimental data and network models will be made available upon publication.

## Methods

### Dataset

Analyses were performed on a previously described dataset of 1-photon calcium imaging (20 Hz sampling frequency) acquired during an olfaction-guided 2-choice decision task^17^. The data are accessible from Zenodo: https://doi.org/10.5281/zenodo.10528244. We considered all mice for which at least 100 neurons were recorded simultaneously, leading to 9 animals of the dataset being eligible for our analysis (total number of neurons across mice: 1949, **Table 1**). Per animal, we used the first available behavioural session with expert performance (mean correct trials: 85.0 ± 1.4%, range: 76.3-89.6%).

**Table 1:** Dataset properties.

| mouse ID | file | session ID | % correct trials | number of cells |
| --- | --- | --- | --- | --- |
| 44 | gcamp6f044_learned.hdf5 | 0 | 84 | 215 |
| 94 | gcamp6f094_learned.hdf5 | 0 | 85 | 133 |
| 216 | gcamp6f216_learned.hdf5 | 0 | 76 | 293 |
| 218 | gcamp6f218_learned.hdf5 | 0 | 82 | 137 |
| 219 | gcamp6f219_learned.hdf5 | 0 | 89 | 229 |
| 478 | gcamp6f478_learning.hdf5 | 2 | 89 | 218 |
| 481 | gcamp6f481_learning.hdf5 | 2 | 85 | 226 |
| 483 | gcamp6f483_learning.hdf5 | 2 | 89 | 230 |
| 485 | gcamp6f485_learning.hdf5 | 2 | 85 | 268 |

### Sequence identification

We initially observed sequential activity using the RasterMap algorithm^22^ (parameters: n_PCs: 200, n_clusters: 100, locality:1, time_lag_window: 20; **Supplementary Fig. 1**), which inspired the following formal analysis. The beginning of odour sampling was determined from 2D tracks of the animals as the frame at which the animal first approached the odour sampling port. Templates were constructed as the distribution of the time to each neuron’s maximal activity in a given trial *T* relative to odour sampling. To avoid spurious influences from weakly active neurons, we only considered cells for which the peak calcium amplitude during the template exceeded 2*SD of the neuron’s activity during the current template time period. We fixed the template duration to 5 s based on the average duration of trials from sampling to reward consumption and on an iterative procedure determining the optimal sequence length (**Supplementary Fig. 1B,C**). To obtain a statistical threshold for sequence reactivation, the calcium activity of each neuron in the template trial was randomly and independently time-shifted (between 2 and 200 seconds, 500 iterations). Then, the mean absolute error between neuron indices in the actual template and the surrogate distribution was measured. The threshold for significant reactivation was defined as the 0.002th percentile of this distribution. A significant match was scored for each time point at which the mean absolute error of the time-to-peak distribution fell below the significance criterion. The trial used for template construction was then moved to trial *T*+1 and the process repeated until all trials had served as the template trial. Reactivation proportion *R* was assessed as

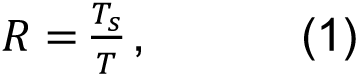

where *T_s_* gives the number of trials with significant matches. This metric therefore measures how many reactivations are detected during the session relative to the total number of trials. Template-match similarity was determined as Spearman’s r between the distribution of indices in the template and the match trial, averaged over all template-match pairings in the session. This metric therefore quantifies to what extent the order of activity peaks remained consistent between template and match.

### Task-phase analysis

For each trial, entry into the sampling zone was defined as task phase 0 and the time point of entry into the reward zone plus 500 ms was defined as task phase 1. The calcium activity of all neurons (i.e., irrespective of whether or not they had contributed to a template in the sequence identification analysis) were expressed as a function of task phase, independently for each trial. We obtained these task-phase tuning functions separately for left- and right-going trials, and separately for the first and second half of the session. Similarity was defined as Pearson’s r between task-phase tuning functions for left- and right-going trials, consistency as Pearson’s r between tuning functions of the first and second half of the session computed separately for both trial types.

### Decoding analysis

To equalize conditions across mice, 100 neurons were randomly selected from each mouse. The decoding results of 100 iterations were averaged. If not indicated otherwise, predictions were made with 5-fold cross-validation with shuffling.

#### Decoding elapsed time from sampling/reward onset

For each trial, we trained a linear support vector machine (sklearn.SVC with linear kernel) on a frame-by-frame basis to predict the current time bin in a time window of 1 s. Trial number was normalized for left- and right-going trials. Decoding performance was expressed as the absolute mean error between predicted and true time. As a control condition, the same predictions were performed with randomly permuted frame times.

#### Decoding task phase

Task phase was predicted by a linear regression model (sklearn.linear_model.LinearRegression with parameter tol=e-8). Decoding performance was expressed as the absolute mean error between predicted and true task phase. For decoding across trial types, the model was trained on all left-going trials and tested on all right-going trials and vice versa, with results expressed as averages across both directions. As a control condition, the same predictions were performed with randomly permuted task phases.

#### Decoding trial outcome from sequential information

For each trial, the time of peak relative to entry into the odour sampling zone or reward zone was measured for each neuron. A logistic regression model (sklearn.LogisticRegression with parameters C=3, max_iter=1e9, tol=1e-13) was trained using time-to-peak and trial type as inputs and the accuracy score as prediction metric. The analysis was run independently for 1, 2 and 4s decoding windows. As a control condition, the same predictions were performed with randomly permuted trial labels.

### Network modelling

#### Task & construction of synthetic target trajectories

The network is trained to learn and reproduce *K* smooth two-dimensional target trajectories generated synthetically. Each trajectory is assembled from three concatenated phases that mirror the task structure: It begins with a stationary period at the origin, corresponding to the sampling epoch. The trajectory then transitions into a condition-specific curved path that represents the movement epoch, with different conditions producing distinct directions of movement. Finally, the trajectory ends with a second stationary period at its endpoint, corresponding to the reward epoch, see **Fig 3B-C**. The trajectories are arranged symmetrically around the vertical axis, with a constant inter-trajectory separation of 20° angle and a total angular spread *Δ* = 20°(*K* − 1). To introduce trial-to-trial variability, we generated independent jittered copies of each prototype trajectory by adding uniform noise with an amplitude of 5% of the maximum coordinate magnitude to the target coordinates. Subsequently, trajectories were smoothed along the time axis with a Gaussian kernel to reduce kinematic discontinuities. Target trajectories were identical between control and penalized runs, so any difference in learning cannot arise from the data. The condition was signaled to the network by a single scalar input *c_k_* ∈ {*k* = 0,…, *K* − 1}, held constant throughout the trial and delivered at every recurrent step.

#### Network architecture and dynamics

The network is a discrete-time recurrent rate network consisting of *N* = 200 units and trained in a closed-loop configuration. At each time step, the network receives an augmented input, 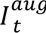, formed by concatenating the static cue *c_k_* with the network’s previously predicted coordinate history. Future positions are zero-filled, and the newly generated output (*x̂_t_*_−1_, *ŷ_t_*_−1_) is written back into 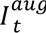 at each step:

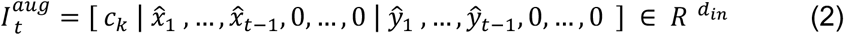

with *d_in_* = 2 × *T* − 1. This creates a closed autoregressive training loop in which the network’s own predictions, rather than the ground-truth targets, drive the recurrence. No teacher forcing or scheduled sampling was applied at any stage of training, the rollout was free-running from the first gradient step to the last, and the loss was computed on the full free-running trajectory. The recurrent layer is composed of 200 sparsely interconnected units with a predefined connectivity determined by a binary mask *M* ∈ {0, 1} *^N^* ^×^ *^N^* that specifies active synaptic connections. *M_ij_* = 1 indicates the presence of a synaptic connection between units (*i*) and (*j*), while *M_ij_* = 0 indicates absence of connection. The effective recurrent weight matrix is formed by element-wise masking of the underlying weight matrix *W_rec_* = *M* ʘ *W̃_rec_*. For each random seed, the mask is sampled once from a Bernoulli distribution (*M_ij_* ∼ *Bernoulli*(*p*)), with (*p*(*M_ij_* = 1) = 0.4), corresponding to approximately 40% of all possible recurrent connections, and remains fixed during training. Consequently, gradients propagate only through the active connections defined by the mask, such that only the existing connections participate in learning.

The hidden state ℎ*_t_* ∈ *R ^N^* updates as:

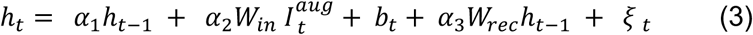

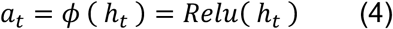

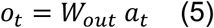

With *a_t_* ∈ *R ^N^* being the neuronal activations, *φ* = *ReLU* the activation function, and the readout *o_t_* ∈ *R ^d_out_^* (*d_out_* = 2) being a bias-free linear projection of the post-activation onto the behavioral plane. All weights, {*W_in_* ∈ *R ^N^* ^×^ *^d_in_^*, *W_rec_* ∈ *R ^N^* ^×^ *^N^*, *W_out_* ∈ *R ^d_out_^* ^× *N*^, *b_t_* ∈ *R ^N^*} were initialized from a uniform variance-scaling distribution. The initial hidden state was sampled from a uniform distribution ℎ_0_ ∼ *U*(0, 1), while per-step state disturbances were sampled from a Gaussian distribution *ξ _t_* ∼ *N*(0, *σ*^2^*I*). All hyperparameters of the closed-loop network are listed in Table 2.

**Table 2:**
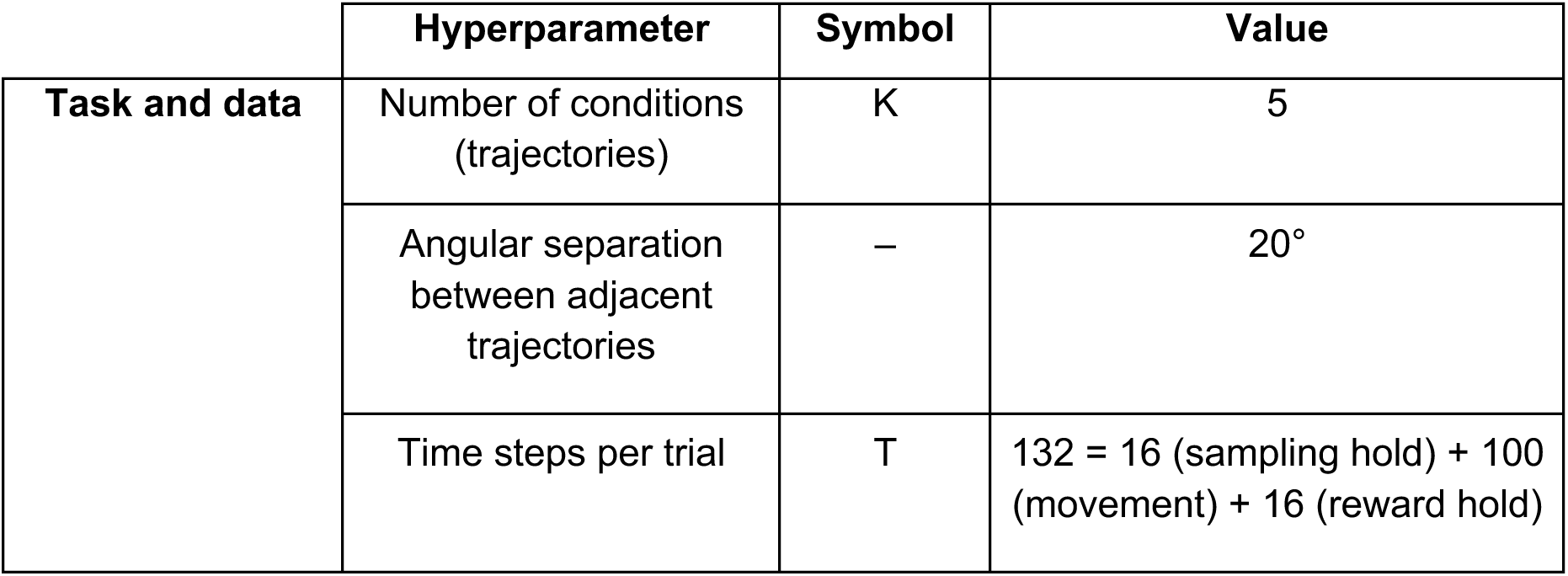
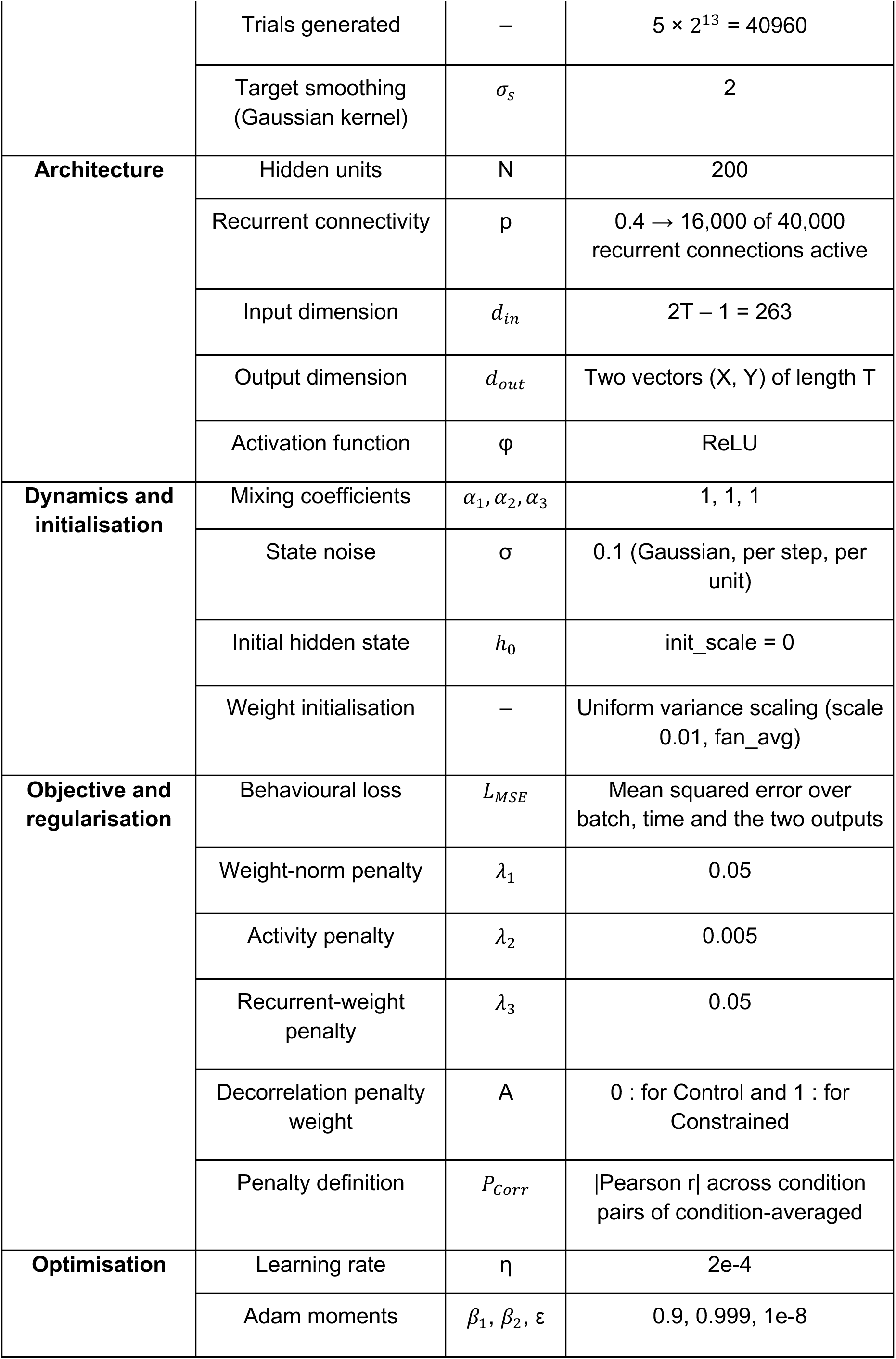
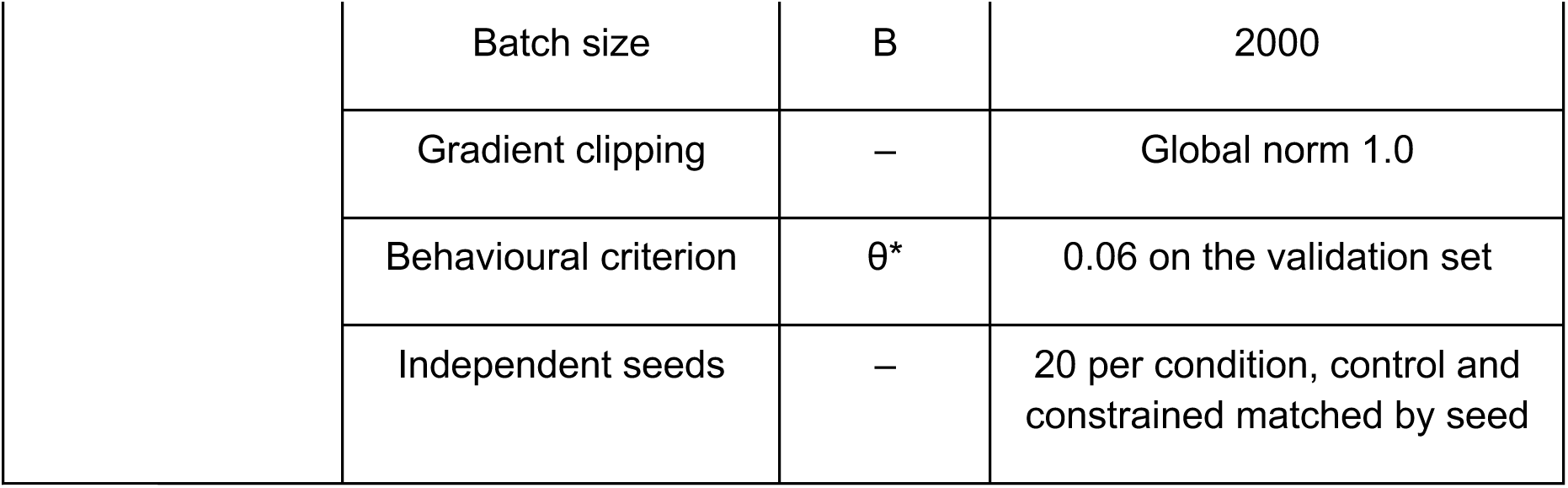
Hyperparameters of the closed-loop recurrent network (Figs. 3–4). Control and constrained networks used identical values; only the penalty weight α differed (0 and 1).

#### Tuning curves and the cross-condition correlation penalty

To manipulate representational sharing, we used a differentiable penalty on the cross-condition similarity of per-unit temporal tuning. Trials were first grouped into *K* categories based on their cue value. The hidden activity of each unit was then averaged across all trials in the same category, producing the tuning curve 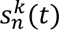 for unit *n* under condition *k*. Each tuning curve was mean-centered over time and normalized, making the penalty invariant to per-unit gain. For each pair of conditions, we computed the Pearson correlation between 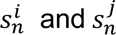 across time for each unit and then averaged the absolute correlation coefficients across units:

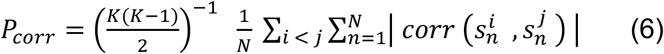

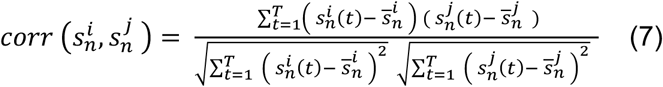

We added *P_corr_* to the objective with weight *α* ≥ 0. Control networks are obtained by setting *α* = 0, while leaving all other aspects of the training pipeline unchanged, so that control and constrained networks differ only in whether *P_corr_* contributes to the gradient. During training, the constrained networks try to minimize this penalty, which drives the temporal activity of each unit to be different or uncorrelated across different conditions, while leaving overall weight statistics unconstrained, see **Supplementary Fig 4**.

#### Objective, optimization and gradient-scale control

The training objective is a composite loss consisting of a behavioral reconstruction term, a regularizer, and the correlation penalty. Networks were trained to minimize the following objective:

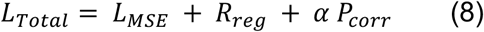

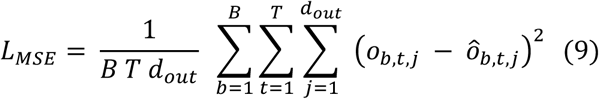

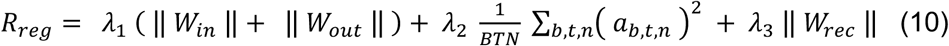

*L_MSE_* is the reconstruction term, defined as the mean-squared error (MSE) between predicted and target trajectories, averaged over batch, time, and the two output dimensions. The resulting *L_MSE_* value served as the behavioral performance metric. Training is considered successful when this term decreases below a predefined loss threshold *θ*\*. The regularization objective *R_re_*_g_ consists of three components. The first constrains the magnitudes of the input and output weight matrices through their Frobenius norms, preventing excessive input and output gains. The second penalizes the mean squared post-activation activity, encouraging sparse and energy-efficient hidden-unit responses. The third constrains the Frobenius norm of the masked recurrent weight matrix, preventing excessively strong recurrent connections while promoting efficient use of the predefined sparse connectivity. For constrained networks, we add the weighted generalization penalty *P_corr_*, which biases the network toward condition-specific representations by penalizing shared sequential structure across conditions.

For control and constrained networks, *α* was set to 0 and 1, respectively. While in both cases *λ*₁, *λ*₂ and *λ*₃ were held constant. The coefficients *α_i_* that govern the contribution of different terms to the network’s hidden activations, the regularization penalty strengths *λ_i_*, and the constraint penalty coefficient *α* used in the constrained network are all fixed hyperparameters during the training process (Table 2). To minimize the training loss and improve task learning, the network parameters are updated using the Adam optimization algorithm^40^, with default hyperparameters. To prevent exploding gradients and improve training stability, gradients were clipped to a maximum global norm before each parameter update. Due to the autoregressive nature of the network, predictions were recursively fed back as inputs during training, resulting in a closed-loop training regime. Unlike teacher forcing, where the ground-truth output from the previous time step is used as the next input, we trained the network in a free-running setting, where each prediction was generated solely from the network’s own previous outputs.

*Activity-constrained control networks*.

To dissociate overall activity from representational sharing, we trained a separate set of seed-matched networks in which *P_act_* was replaced by a penalty on the population activity level. Activity was quantified as the root-mean-square post-activation activity over batch, time and units:

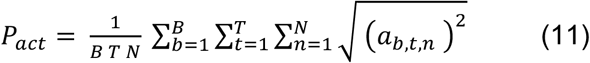

and penalized relative to a target A*:

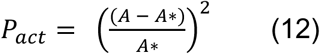

For each seed, A* was set to 0.5 × the activity of the seed-matched control network, measured on a held-out batch after training. Because the activity term of the regularizer (λ₂, Objective section) also acts on hidden-unit activity and would bias networks toward zero activity, competing with the imposed target, it was disabled (λ₂ = 0) for both control and activity-constrained networks in this manipulation, while λ₁ and λ₃ were kept unchanged. The activity penalty was added to the objective with α = 1 (*L_Total_* = *L_MSE_* + *R_re_*_g_ + *α P_act_*). All other hyperparameters, initialization, connectivity mask and batch ordering were identical between the two networks of each seed-matched pair. Training terminated when the running minimum of the task loss fell below θ* while A lay within ±5% of A*, ensuring that activity-constrained networks were timed only once they solved the task at the imposed activity level.

#### Quantification and statistical analysis of learning efficiency

Modelling results are reported across independent seeds. For each seed, the recurrent weights, connectivity mask, dynamical noise, and data ordering were independently re-initialized. Control and constrained networks were matched seed by seed, such that the two architectures differed only in the presence of the decorrelation penalty and all comparisons were paired. Learning efficiency is quantified as the number of optimization steps required to reach a fixed level of behavioural performance. Throughout training we logged the task loss (MSE between predicted and target trajectories) separately from the decorrelation penalty, and all criteria below were applied to the task loss alone, so that control and constrained networks were timed against an identical behavioural measure. Training was terminated at the first epoch in which the validation MSE (task loss only) fell below *θ*^∗^ = 0.06, or at a ceiling of 200,000 epochs. The number of epochs taken to reach this criterion, E*, served as a single summary measure of the time required to reach a fixed level of behavioural performance. We chose the loss *θ*^∗^ = 0.06 as the early stopping loss value because, beyond this point, further training cannot meaningfully improve the fit. Both learning curves have flattened out by this stage. Networks were trained in matched pairs. For each seed, the control and constrained network shared the same initialization, sparse connectivity mask and batch ordering, and differed only in whether the decorrelation penalty contributed to the gradient. The learning-efficiency analysis used *n* = 20 seeds per condition, with trajectories separated by 20°, and was run for *K* = 5.

To assess whether the learning-speed difference depended on the choice of convergence criterion, we repeated the epochs-to-criterion analysis across a ladder of 45 loss criteria spanning *θ* ∈ [0.06,…, 0.5] in increments of 0.01. For each network and criterion, *E*(*θ*) was defined as the first training step at which the logged task loss fell to or below *θ*. This procedure yielded an epochs-to-criterion curve for each network, summarized across seeds. The analysis was performed on control and constrained networks trained on the *K* = 5 task. Networks were matched seed-by-seed, yielding a fully within-subjects design. We fitted a two-way repeated-measures ANOVA with Network (control vs. constrained) and Loss criterion (45 levels) as within-subject factors, with seed as the repeated-measures unit. The Network × Criterion interaction was used to assess whether the difference in learning speed between networks varied as the convergence criterion became more stringent. To determine at which criteria the two networks differed, we performed post-hoc paired t-tests between control and constrained networks at each of the 45 criteria, exploiting the seed-matched design. The resulting *p*-values were Bonferroni-corrected across the 45 comparisons and evaluated at *α_bonf_* = 0.05.

#### Recurrent neural network model

To test whether the sequences depend on the closed-loop architecture, we trained a vanilla recurrent network that receives the cue as external input and has no access to its own output (**Supplementary Fig. 7**). The network had N = 200 tanh units and three input channels: a fixation signal and the cosine and sine of the target direction. A linear readout predicts the (x, y) position at each of the T = 132 time steps. The hidden state was integrated with a leak rate β = 0.2:

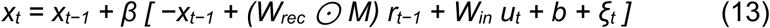

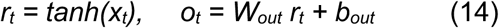

The recurrent connectivity was constrained by a binary mask (M), with 80% of connections retained (Bernoulli, p = 0.8). The mask was sampled independently for each random seed and held fixed throughout training. State noise, *ξ_t_ ∼ N(0, 0.2*^2^), was applied and initial states were sampled from *U(−0.1, 0.1*), weights were initialized from *N(0, 0.1*^2^), and biases were initialized to zero.

#### Model training and performance evaluation

For this model, control (α = 0) and constrained (α = 1) networks were trained as 10 seed-matched pairs with identical initial weights, mask and data split. Control networks were trained to minimize the mean squared error across the entire trial, including the initial hold period. Constrained networks were trained using an augmented objective, (L = L^MSE^ + α P^corr^), where Pcorr is the cross-condition correlation penalty on per-unit tuning curves defined for the closed-loop network (see ‘Tuning curves and the cross-condition correlation penalty’): the absolute Pearson correlation between a unit’s condition-averaged time courses for each pair of conditions, averaged over units and summed over all pairs of conditions. Networks were optimized using Adam with a learning rate of 1e-3 and global gradient norm clipping at 0.2. Training was terminated when the validation MSE met the same predefined stopping criterion or when the maximum number of epochs was reached. The number of epochs required to meet this criterion was recorded as the epochs-to-criterion.

### Statistics and software packages

Comparisons between two independent groups were performed with unpaired Student’s t-tests whenever the data were normally distributed (tested with a Shapiro-Wilk test). Non-normal data were compared with a Mann-Whitney U-test. Dependent two-group data were assessed with paired t-tests or Wilcoxon signed-rank tests, depending on normality. Individual time series were assessed with a one-way repeated measures ANOVA. Multi-timepoint statistics for two data series were obtained with a two-way repeated measures ANOVA followed by post-hoc testing with paired t-tests or Wilcoxon signed-rank test with Šidák correction. All tests used a two-sided design. Experimental data are based on n=9 animals, modelling results on n=20 independent seeds unless indicated otherwise. Statistical tests were computed in python using *scipy.stats* and *pingouin*^41^ packages.

Recurrent network models were implemented in TensorFlow/Keras, with array computations in NumPy and additional numerical routines from SciPy. Dimensionality reduction and decoding analyses used scikit-learn, and data handling and visualization used pandas and Matplotlib.

### Use of generative AI

Large language models (LLMs) were used for language editing in this work. LLMs were moreover used to assist with implementation details of the code used for data analysis and network modelling (e.g., debugging, documentation of statistical tests). All experimental designs, network architectures, statistical analyses, and interpretation of results were conceived and carried out by the authors.

## Supplementary Figures

**Supplementary Fig. 1:**
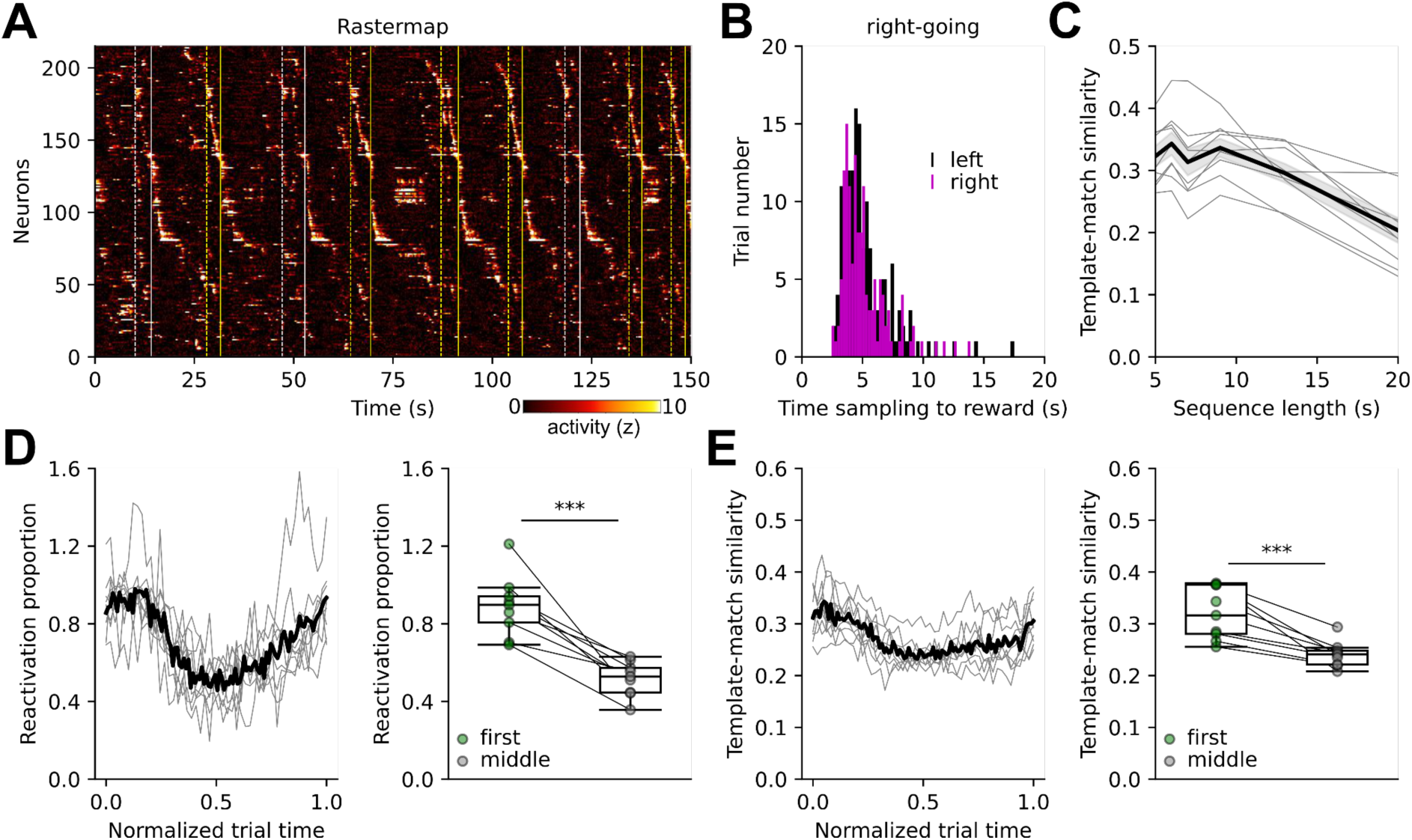
Neuronal sequence identification. **A** Rastermap visualisation of neuronal sequences from an example mouse. Dashed lines indicate entry into the sampling zone, solid lines show entries into the reward zone. **B** Histogram of elapsed time of left-going (black) and right-going trials (magenta) from entry to the sampling zone to entry to the reward Zone (i.e., trial duration) across all mice. Average trial duration does not differ for left- and right-going trials. p=0.82, W=20, Wilcoxon signed-rank test. **C** Template-match similarity for different sequence lengths used for template matching ranging from 5 to 20 s. Optimal similarity is achieved for timescales that match actual trial durations but decreases for longer time intervals. p=4.6e-9, F=17.2, one-way repeated measures ANOVA. For all analysis, a sequence length of 5 s was chosen since it falls in the plateau of optimal template-match similarities. **D** Template-match similarity for random template start points in normalized trial time (0: entry into sampling phase, 1: return to sampling phase). Right: Comparison of reactivation proportion for normalized trial time zero (first) vs. normalized trial time 0.5 (middle). p=0.0003, t=6.1, paired t-test. **E** Same as D but for template-match similarity. p=0.0008, t=5.2, paired t-test. All data based on n=9 mice. Circles in D, E show mouse averages, boxes show median and first and third quartiles. Thick lines in C, D, E show mean, grey lines individual mice, shaded area sem.

**Supplementary Fig. 2:**
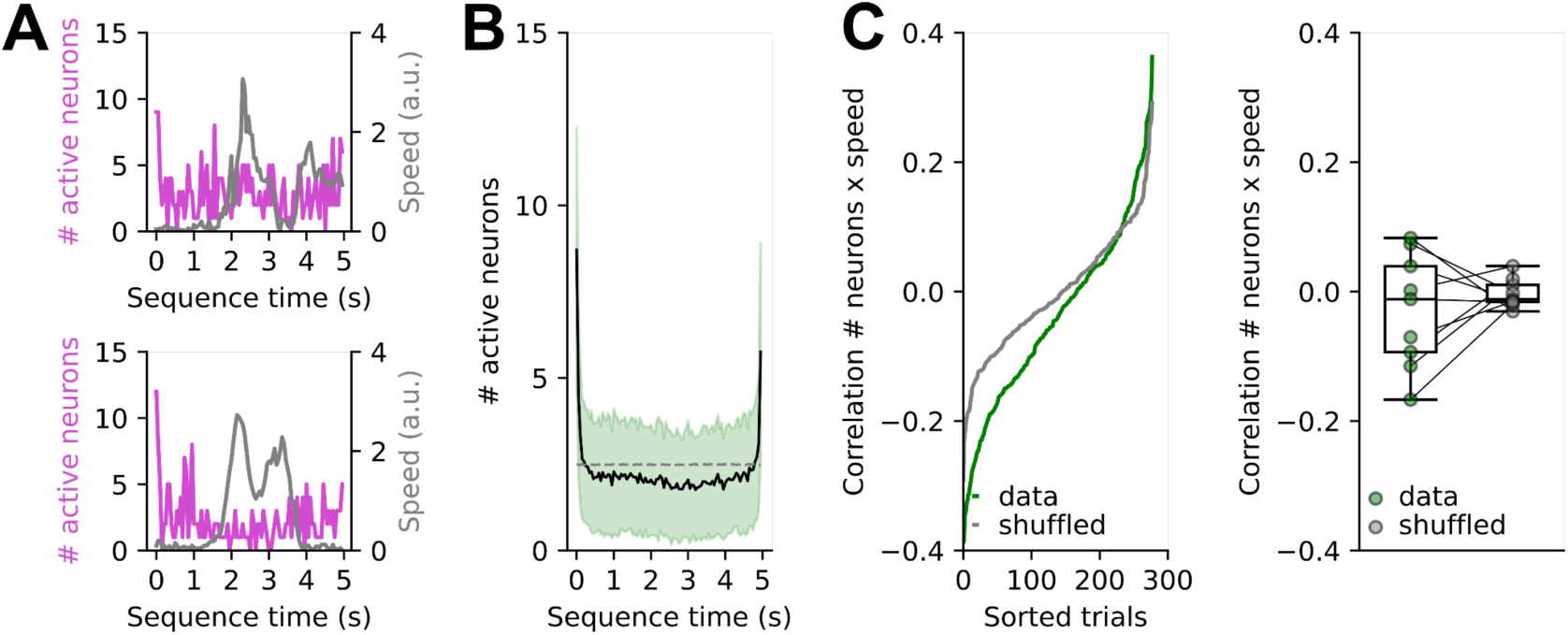
Sequence contribution during immobile and movement epochs. **A** Two examples of single template trials. Shown are the number of neurons active as a function of sequence progression time (template length: 5 s, with zero referring to the entry into the sampling zone, magenta) and the speed profile of the trial. **B** Number of active neurons as a function of sequence progression time (black: average, green area: SD, n=278 templates). The dashed gray line indicates the 99^th^ percentile of a null distribution (permuted neuron numbers, 5000 iterations), which is exceeded by the data mean only at <0.25 s and >4.75 s. **C** Left: Correlation of neuron number and speed (i.e., the vectors shown in magenta and gray in A, for all templates across mice (n=278 templates). Shuffled data were obtained by randomly permuting the speed profile of each trial. Right: Average correlation per mouse does not differ from shuffled data, arguing against an influence of the animals current speed on sequence participation. p=0.428, t=-0.83, paired t-test. All data based on n=9 mice. Circles in C show mouse averages, boxes show median and first and third quartiles. Thick line in B shows the mean, shaded area SD.

**Supplementary Fig. 3:**
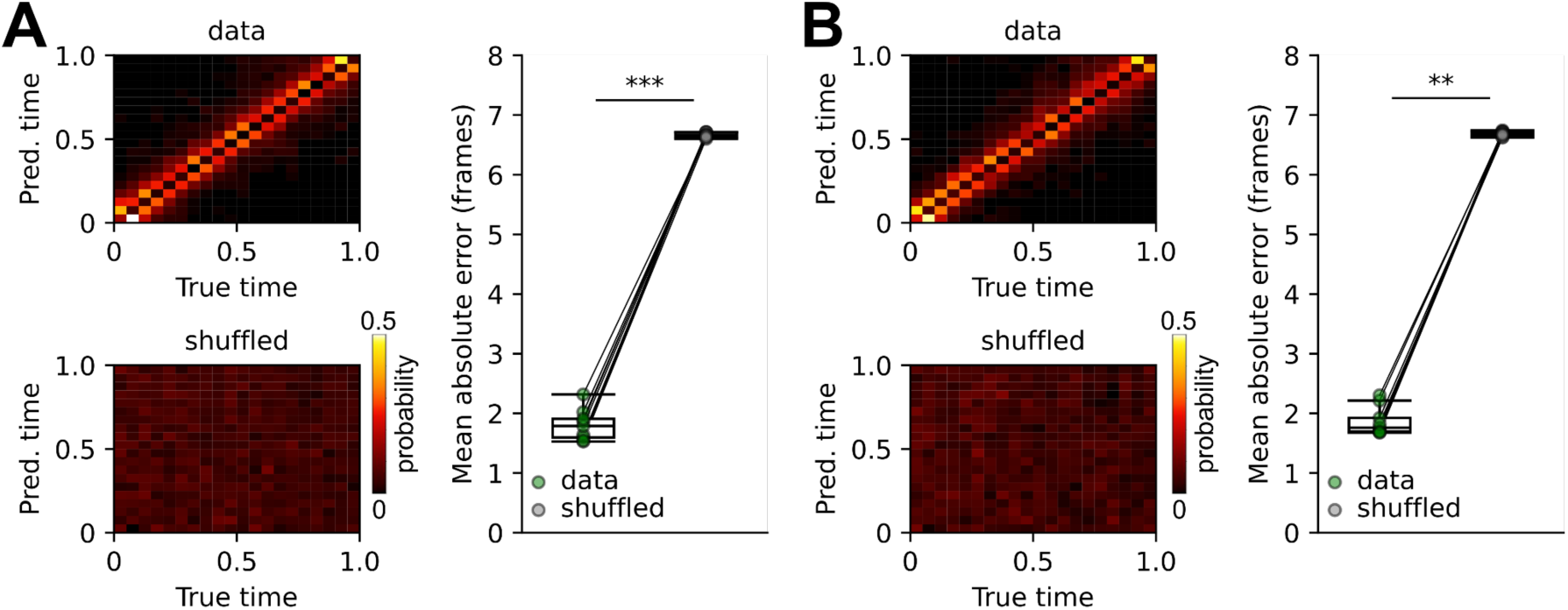
Decoding elapsed time during immobile task epochs. **A** Decoding elapsed time from entry into the sampling zone (timepoint 0). Left: Confusion matrices show decoding results of a linear SVC trained on data (top) and on shuffled time frames (bottom). Right: Average decoding error is lower for data-trained models (p=e-11, t=-52.6, paired t-test). **B** Same as A but for the time from entry into the reward zone. p=0.003, W=0, Wilcoxon signed-rank test). All data based on n=9 mice with 100 randomly selected neurons each, averaged over 100 iterations. Circles show mouse averages, boxes show median and first and third quartiles.

**Supplementary Fig. 4:**
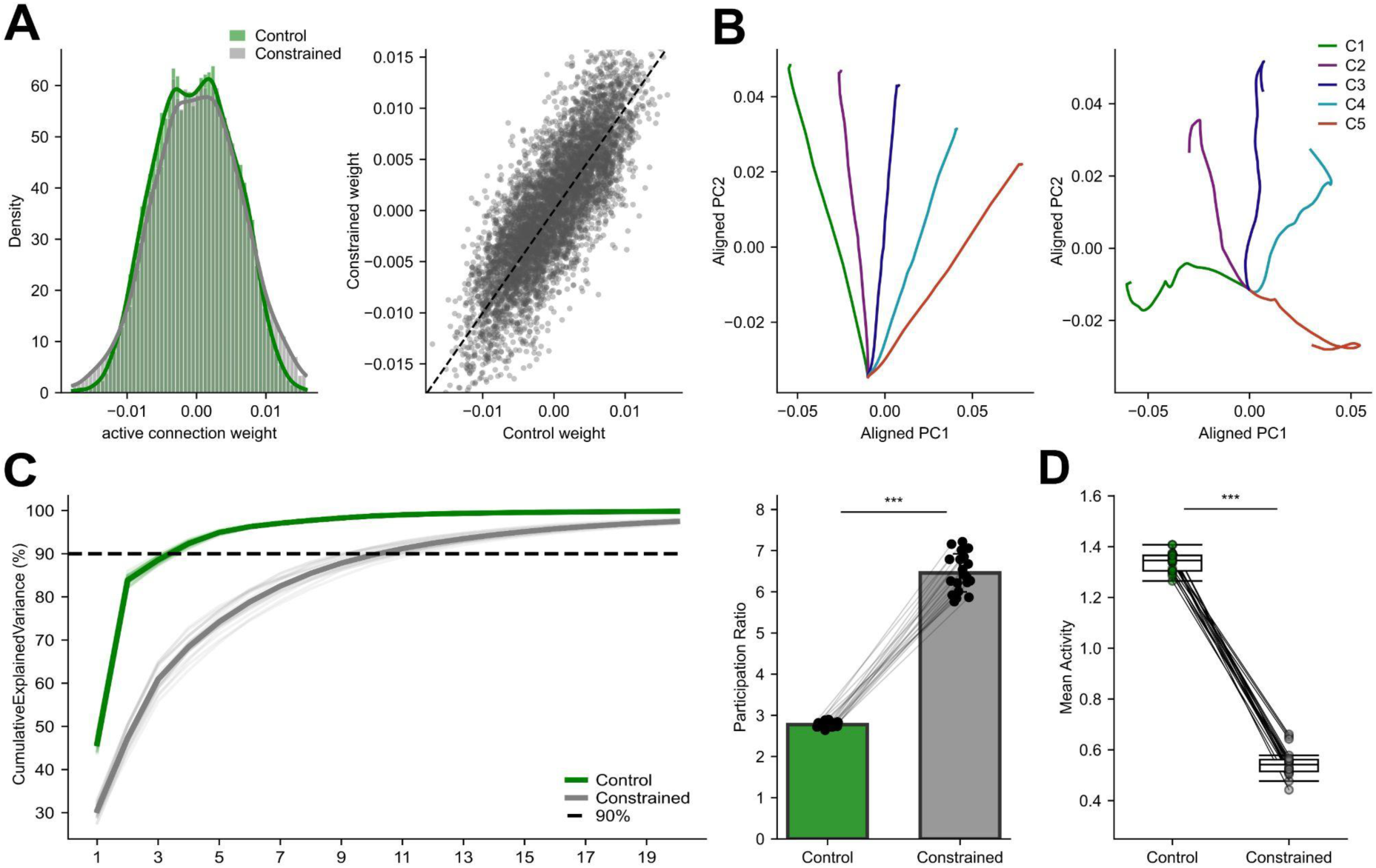
The correlation penalty reorganizes population geometry without altering network connectivity. **A** Preservation of recurrent connection-weight magnitudes under the constraint. **Left**: distribution of active (unmasked) recurrent connection weights for a representative control and constrained network; the two distributions are nearly indistinguishable in shape and statistically equivalent within a pre-specified margin of ±0.00063 (10% of the pooled SD; two one-sided tests, dof = 15,999, TOST p = 2.02e-20). **Right**: per-connection comparison of weight magnitude between a constrained network and its seed-matched control (dashed line: identity); weights are strongly correlated (Pearson r = 0.799) and cluster along the identity line. **B** Two-dimensional PCA projections of the population trajectories for all five conditions in a representative control network (left) and its seed-matched constrained network (right, α = 1). In the control network, all five trajectories leave a common origin and travel along similar, closely aligned paths, consistent with the geometric signature of a shared low-dimensional scaffold. In the constrained network, trajectories leave the same origin but follow distinct, condition-specific routes with their own curvature, consistent with the higher-dimensional scaffold; constrained trajectories diverge into distinct, condition-specific routes. **C** Increased dimensionality in constrained networks. Left: cumulative variance explained by successive principal components of trial-averaged hidden-unit activity for control and constrained networks (n = 20 seeds each); dashed line marks the 90% threshold. Right: participation ratio for control versus constrained networks; constrained networks occupy a significantly higher-dimensional subspace than controls (paired t-test, |t| = 34.68, p = 1.21e-18, dof = 19, n = 20 seeds). **D** Constrained networks show significantly reduced mean hidden-unit activity relative to their seed-matched controls, averaged across all five conditions (control: 1.340 ± 0.038; constrained: 0.547 ± 0.054, mean ± SD; paired t-test, t = 51.39, p = 7.4e-22, dof = 19, n = 20 seeds).

**Supplementary Fig. 5:**
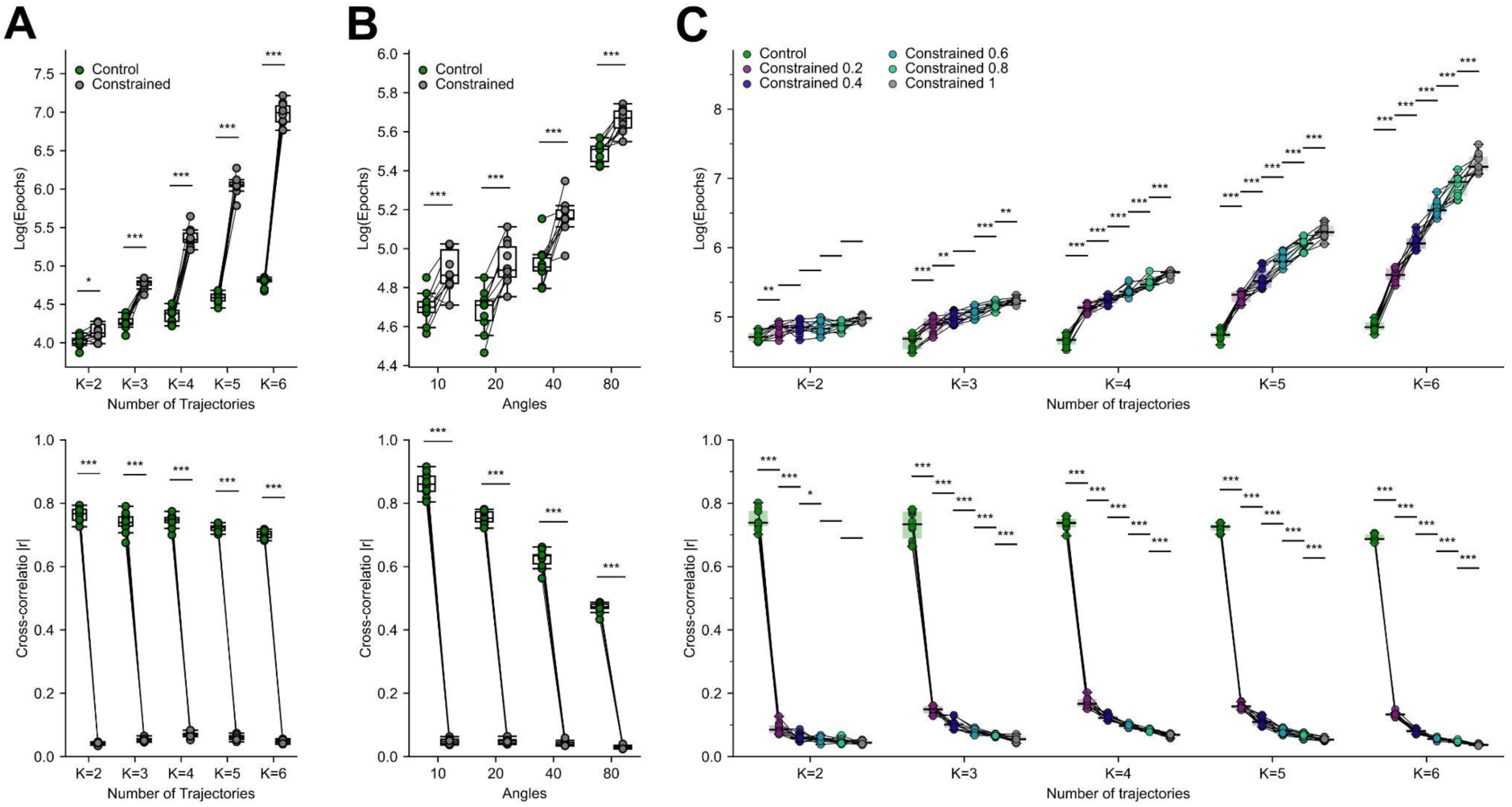
The learning cost of losing shared task-progress coding is robust to dataset size, trajectory geometry, and scales with penalty strength. **A** Dataset size does not account for the training gap. Top: Epochs required to reach criterion performance for control and constrained networks trained on an increasing number of trajectory conditions, K ranging from 2 to 6, with total training data held constant. Constrained networks required significantly more epochs than controls at every K (paired t-test, dof = 9: K=2, t = −2.70, p = 0.024; K=3, t = −14.95, p = 1.16e-7; K=4, t = −16.95, p = 3.88e-8; K=5, t = −20.61, p = 6.95e-9; K=6, t = −18.57, p = 1.75e-8). Bottom: Correlation between condition-averaged tuning curves in control and constrained networks. Despite equalized dataset size, constrained networks exhibited significantly and near-zero cross-condition correlation relative to controls at every K (paired t-test, dof = 9: K=2, t = 119.22, p = 1.04e-15; K=3, t = 60.43, p = 4.69e-13; K=4, t = 89.99, p = 1.31e-14; K=5, t = 124.10, p = 7.28e-16; K=6, t = 146.00, p = 1.69e-16, n=10), and constrained networks required progressively more epochs as K increased, whereas control networks remained fast throughout. These results rule out training-data quantity as an explanation for the widening training gap observed in the trajectory-number experiment (**Fig. 4H**). **B** Effect of angular separation between trajectory conditions. Top: Epochs required to reach criterion performance for control and constrained networks with the number of trajectory conditions fixed at K=2, while the angular separation between the two target trajectories is varied (10°–80°). Epochs-to-criterion increased with angular separation for both network types, with constrained networks consistently requiring significantly more epochs than control networks at every angle tested (paired t-test, dof = 9: 10°, t = −5.66, p = 3.11e-4; 20°, t = −6.87, p = 7.28e-5; 40°, t = −5.32, p = 4.81e-4; 80°, t = −7.00, p = 6.33e-5). Bottom: Correlation between condition-averaged tuning curves. Correlation in control networks declined modestly as angular separation increased, whereas constrained networks remained near zero across all angles tested, with control networks showing significantly higher cross-condition correlation than constrained networks at every angle (paired t-test, dof = 9: 10°, t = 71.29, p = 1.06e-13; 20°, t = 102.22, p = 4.16e-15; 40°, t = 61.91, p = 3.78e-13; 80°, t = 82.11, p = 2.98e-14). **C** Learning cost scales with constraint strength. **Top**: Epochs required to reach criterion performance (log scale) as a function of the number of trajectory conditions (K) for control networks and constrained networks trained with penalty weights α = 0.2, 0.4, 0.6, 0.8, and 1. Epochs-to-criterion increased monotonically with both K and penalty dose (one-way repeated-measures ANOVA across all six network conditions, run separately at each K: K=2, F = 25.41, p = 3.48e-16; K=3, F = 125.36, p = 3.05e-40; K=4, F = 379.07, p = 4.17e-61; K=5, F = 377.60, p = 4.97e-61; K=6, F = 300.52, p = 1.40e-56). Bottom: Correlation between condition-averaged tuning curves as a function of K for the same networks. Correlation dropped sharply upon introducing the penalty, even at the lowest dose (α = 0.2), and decreased further with increasing penalty dose (one-way repeated-measures ANOVA at each K: K=2, F = 8212.11, p = 1.46e-123; K=3, F = 4460.74, p = 5.15e-111; K=4, F = 13503.79, p = 8.37e-134; K=5, F = 16476.78, p = 6.66e-138; K=6, F = 23896.65, p = 1.45e-145). Pairwise comparisons between adjacent penalty doses (brackets) are Bonferroni-corrected paired t-tests.

**Supplementary Fig. 6:**
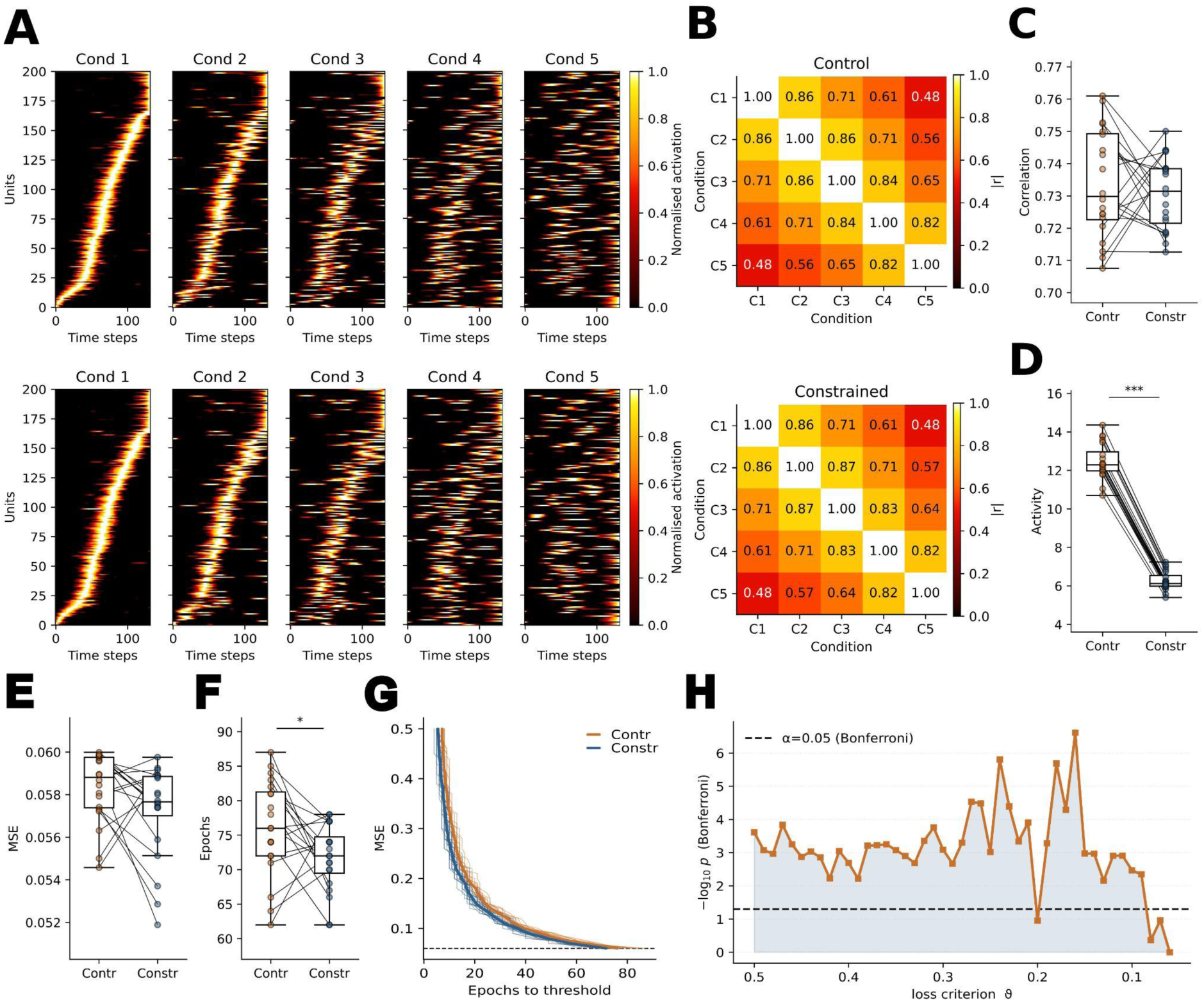
Reducing overall activity preserves generalized task-progress coding and learning efficiency. **A** Trial-averaged hidden-unit activation heatmaps for each of the five conditions in a representative control network (top) and its seed-matched activity-constrained network (bottom; A* = 0.5 × control activity). Units are sorted by peak-activation latency, with the ordering fixed from Condition 1 (as in Fig. 3E). Unlike the correlation penalty (Fig. 4A), the activity penalty preserves the diagonal band of sequential activity shared across conditions. **B** Representational similarity matrices (mean absolute Pearson correlation, |r|, between condition-averaged hidden-unit tuning curves) for the networks in A. Cross-condition similarity is virtually identical in both networks. **C** Mean cross-condition |r| for control and activity-constrained networks (control: 0.734 ± 0.016; activity-constrained: 0.731 ± 0.011, mean ± SD; n = 20 seeds). Cross-condition similarity did not differ between networks (paired t-test, t = 0.61, p = 0.547, dof = 19) and was statistically equivalent (TOST equivalence test, paired, bound = ±0.067 [10% of the reduction in |r| produced by the correlation penalty, Fig. 4B], dof = 19, p = 2.4e-11). **D** Overall hidden-unit activity (root-mean-square post-activation activity) was halved by the activity penalty (control: 12.46 ± 0.91; activity-constrained: 6.26 ± 0.48; paired t-test, t = 62.48, p = 1.85e-23, dof = 19). **E** Final post-convergence task MSE, showing statistically equivalent asymptotic performance (TOST equivalence test, paired, bound = ±0.006 [10% of θ*], dof = 19, p = 1.4e-7). **F** Epochs required to reach the behavioral criterion θ* = 0.06 (control: 76.1 ± 6.8; activity-constrained: 71.7 ± 4.7; paired t-test, t = 2.17, p = 0.043, dof = 19). **G** Task MSE as a function of training epoch for control and activity-constrained networks (thin lines: individual seeds; thick lines: mean), dashed line indicates θ* = 0.06. **H** Bonferroni-corrected significance (−log₁₀ p) of the control vs activity-constrained difference in epochs- to-criterion across 45 loss criteria spanning θ ∈ [0.06, 0.5] (paired t-tests, dof = 19; dashed line, α = 0.05). Activity-constrained networks learned slightly faster across most criteria (two-way repeated-measures ANOVA, main effect of network: F(1,19) = 272.61, p = 1.0e-12). The difference diminished at stricter criteria (network × criterion interaction: F(44,836) = 3.31, p = 3.9e-3) and was not significant at θ* after correction (|t| range: 2.17–9.98; Bonferroni-corrected p range: 2.4e-7 to 1). In contrast to the correlation penalty, reducing activity did not impair learning efficiency.

**Supplementary Fig. 7:**
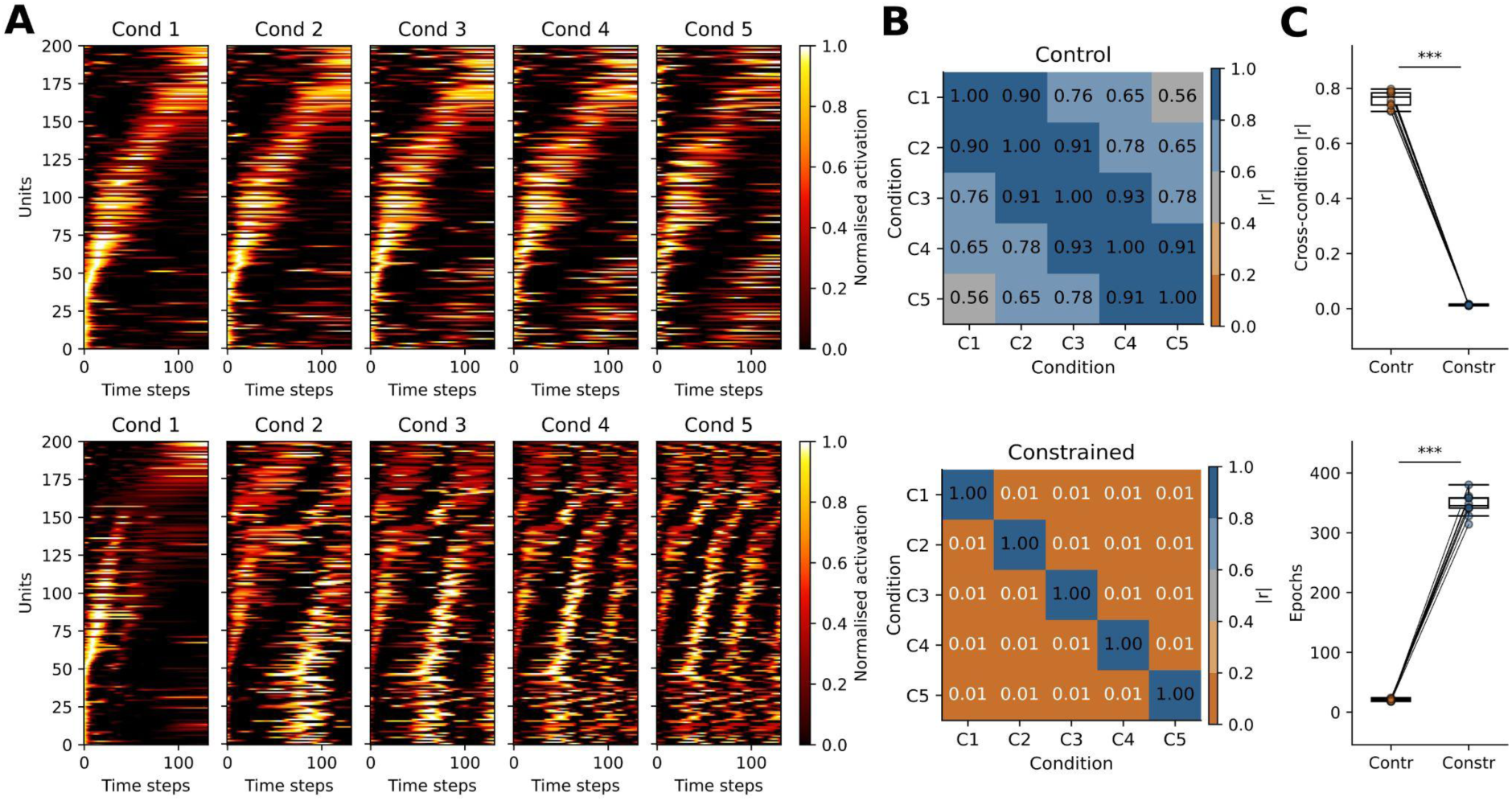
Shared task-progress sequences speed up learning in a vanilla recurrent network, independent of output feedback. The open-loop control was implemented using a vanilla RNN that receives the cue as an external input but does not use its own outputs, in contrast to the closed-loop network shown in Figs. 3–4 (see Methods). The seed-matched control (α = 0) and constrained (α = 1) networks differed only in the decorrelation penalty. **A** Trial-averaged hidden-unit activation heatmaps for each of the five conditions (Cond 1–5) in a representative control network (α = 0, top) and its seed-matched constrained network (α = 1, bottom). Units are sorted by peak-activation latency, with the ordering fixed from Condition 1. In the control network, a diagonal band of sequential activity is visible across all conditions, indicating a shared temporal scaffold. In the constrained network, sequential structure is largely preserved only for the condition used to define the sort order (Cond 1). **B** Representational similarity matrices (mean pairwise absolute Pearson correlation, |r|, between condition-averaged hidden-unit tuning curves) for the networks shown in A. Control networks show strong cross-condition similarity, whereas constrained networks show uniformly low similarity. **C Top**: Mean cross-condition |r| for control and constrained networks (lines connect seed-matched pairs). Constrained networks showed significantly lower cross-condition correlation than controls (control: 0.76 ± 0.03; constrained: 0.013 ± 0.002, mean ± SD; paired t-test, t = 77.37, p = 5.1e-14, dof = 9, n = 10 seeds). **Bottom**: Epochs required to reach a behavioral criterion. Constrained networks required significantly more epochs than seed-matched controls (control: 20.3 ± 2.2; constrained: 347.1 ± 18.5, mean ± SD; paired t-test, t = −54.4, p = 1.2e-12, dof = 9, n = 10 seeds), reproducing the learning cost observed in the closed-loop network (Fig. 4).

